# Functional requirement of the *Arabidopsis* importin-α nuclear transport receptor family in autoimmunity mediated by the NLR protein SNC1

**DOI:** 10.1101/2020.05.09.084020

**Authors:** Daniel Lüdke, Charlotte Roth, Sieglinde A. Kamrad, Jana Messerschmidt, Denise Hartken, Jonas Appel, Bojan F. Hörnich, Qiqi Yan, Stefan Kusch, Melanie Klenke, Annette Gunkel, Lennart Wirthmueller, Marcel Wiermer

**Author notes:** These authors contributed equally to the work (Daniel Lüdke, Charlotte Roth).

## Abstract

IMPORTIN-α3/MOS6 (MODIFIER OF SNC1, 6) is one of nine importin-α isoforms in *Arabidopsis* that recruit nuclear localization signal (NLS)-containing cargo proteins to the nuclear import machinery. *IMP-α3*/*MOS6* is required genetically for full autoimmunity of the nucleotide-binding leucine-rich repeat (*NLR*) immune receptor mutant *snc1* (*suppressor of npr1-1, constitutive 1*) and *MOS6* also contributes to basal disease resistance. Here, we investigated the contribution of the other *importin-*α genes to both types of immune responses, and we analyzed potential interactions of all importin-α isoforms with SNC1. By using reverse-genetic analyses in *Arabidopsis* and protein-protein interaction assays in *N. benthamiana* we provide evidence that among the nine α-importins in *Arabidopsis*, IMP-α3/MOS6 is the main nuclear transport receptor of SNC1, and that *IMP-α3*/*MOS6* is required selectively for autoimmunity of *snc1* and basal resistance to mildly virulent *Pseudomonas syringae* in *Arabidopsis*.

**SIGNIFICANCE STATEMENT:** Specific requirement for the *Arabidopsis* α-importin MOS6 in *snc1*-mediated autoimmunity is explained by selective formation of MOS6-SNC1 nuclear import complexes.

## INTRODUCTION

In eukaryotic cells, the nuclear envelope separates the nucleoplasm from the surrounding cytoplasm. It acts as a protective compartment boundary for the genome, but it also provides eukaryotic cells with an important regulatory feature to control the specificity and timing of signaling pathways and gene expression in response to both cellular and environmental stimuli (Kaffman and O’Shea, 1999; Orphanides and Reinberg, 2002; Gu, 2018). The nuclear envelope consists of an inner and outer lipid bilayer and is spanned by a multitude of nuclear pore complexes (NPCs) that fuse both membranes to form a central channel for the selective bidirectional transport of macromolecules as well as the passive diffusion of small soluble molecules <40-60 kDa between the nucleoplasm and the cytoplasm (Stewart, 2007; Wang and Brattain, 2007; Beck and Hurt, 2017). The approximately 30 different protein constituents of NPCs can have predominantly structural functions or play active roles in nuclear transport and are collectively termed nucleoporins (NUPs). NUPs containing intrinsically disordered domains of hydrophobic phenylalanine (F)-glycine (G) repeats form a meshwork in the central transport channel that is crucial for the function of NPCs (Grossman *et al.*, 2012; Tamura and Hara-Nishimura, 2013; Beck and Hurt, 2017). This meshwork of FG-NUPs creates the selective permeability barrier and provides binding sites for nuclear transport receptors (NTRs) that traverse the central channel of the NPC during facilitated transport (Schmidt and Görlich, 2016). NTRs include both importins and exportins which recognize localization signals on their cargos and mediate nuclear import and export, respectively (Pemberton and Paschal, 2005; Wente and Rout, 2010), albeit NTRs that mediate bidirectional transport have also been described (Mingot *et al.*, 2001).

The transport of proteins destined for active nuclear import is generally mediated by an importin-α and importin-β heterodimer. α-importins function as adapter proteins that recognize and bind to nuclear localization signals (NLSs) of cargo proteins. The best characterized NLSs are comprised of one (monopartite) or two (bipartite) sequence motifs enriched in the basic amino acids lysine (K) and arginine (R). These canonical or classical NLSs (cNLSs) have the consensus sequences [K(K/R)X(K/R)] and [(K/R)(K/R)X_10-12_(K/R)_3/5_], respectively (Chang *et al.*, 2012; Marfori *et al.*, 2012). Importin-α proteins possess a central series of ten armadillo (ARM) repeats that form the NLS-binding cleft with two distinct NLS contact sites on its concave side, referred to as the major and minor NLS binding site. Whereas both sites simultaneously interact with bipartite cNLSs, monopartite cNLSs preferentially bind to the major site (Marfori *et al.*, 2011; Chang *et al.*, 2013; Christie *et al.*, 2016). In addition to the ARM repeat domain, importin-α family members typically contain an N-terminal α-helix that mediates direct contact to importin-β and is therefore termed the importin-β-binding (IBB) domain. The flexible IBB domain has a dual regulatory function. On the cytoplasmic side of the NPC, it links the importin-α/NLS-cargo complex to the importin-β carrier molecule. Importin-β subsequently mediates the active transport of the ternary complex into the nucleus by directly interacting with the FG-NUP meshwork in the central channel of the NPC (Kobe, 1999; Cook *et al.*, 2007; Chang *et al.*, 2012; Marfori *et al.*, 2012). In the nucleoplasm, the ternary import complex is destabilized by binding of importin-β to the small GTPase RAS-RELATED NUCLEAR PROTEIN (RAN) in its GTP-bound form, resulting in dissociation of the importin-α IBB domain from importin-β. The IBB domain harbors a cluster of basic amino acids that is related to bipartite NLSs and competes with the NLSs of cargo proteins for binding to the ARM repeats (Kobe, 1999). This auto-inhibitory effect of the IBB domain helps to free the cargo from importin-α ARM-repeats inside the nucleoplasm after the IBB domain is released from importin-β by RAN·GTP (Moroianu *et al.*, 1996; Kobe, 1999; Harreman *et al.*, 2003; Cook *et al.*, 2007; Chang *et al.*, 2012). Subsequently, the dimeric importin-β/RAN·GTP complex is exported to the cytoplasm, whereas a C-terminal acidic patch of importin-α interacts with the RAN·GTP-bound export carrier CELLULAR APOPTOSIS SUSCEPTIBILITY (CAS) for recycling of cargo-free importin-α back into the cytoplasm (Goldfarb *et al.*, 2004; Christie *et al.*, 2016). In the cytoplasm, RAN GTPase-ACTIVATING PROTEIN (RanGAP) and its co-factor RAN GTP BINDING PROTEIN1 (RanBP1) impart GTP hydrolysis on RAN to release importin-α and importin-β for another round of cargo import (Stewart, 2007).

The *importin-α* gene family has expanded substantially during the course of eukaryotic evolution, probably reflecting adaptation towards a more complex, tissue- and/or stimulus-specific regulation of nuclear protein import (Yano *et al.*, 1992; Yasuhara *et al.*, 2007; Hu *et al.*, 2010; Kelley *et al.*, 2010; Pumroy and Cingolani, 2015). The genome of the model plant species *Arabidopsis thaliana* encodes nine importin-α paralogs, of which the isoforms *α1* - *α4*, *α6* and *α9* are expressed ubiquitously based on publicly available gene expression data (Wirthmueller *et al.*, 2013). *Arabidopsis IMPORTIN-α3* (*IMP-α3*), also termed *MOS6* (for *MODIFIER OF SNC1, 6*), was identified in a forward genetic screen for components that contribute to constitutive defense activation and related growth inhibition of the autoimmune mutant *suppressor of npr1-1, constitutive1* (*snc1*; Li *et al.*, 2001; Palma *et al.*, 2005). Constitutive immune pathway activation in *snc1* plants is caused by an E_552_K mutation in the nucleotide-binding leucine-rich repeat (NLR) protein SNC1 (Zhang *et al.*, 2003). Plant NLR proteins function as intracellular immune receptors that perceive pathogen-secreted virulence factors (effectors) to activate a defense response termed effector-triggered immunity (ETI). ETI is commonly associated with a hypersensitive cell death response (HR) at the infection site to restrict pathogen spread (Cui *et al.*, 2015), albeit auto-activation of SNC1 *per se* does not cause spontaneous cell death in the *snc1* mutant (Li *et al.*, 2001; Zhang *et al.*, 2003). SNC1 is a nucleocytoplasmic Toll-Interleukin Receptor (TIR)-type NLR (TNL) protein and its nuclear pool is essential for the autoimmune phenotype of *snc1* plants (Cheng *et al.*, 2009; Zhu *et al.*, 2010a; Wiermer *et al.*, 2010). Therefore, the identification of three mutant alleles of *mos6* as genetic suppressors of *snc1* autoimmune phenotypes suggest that MOS6 imports the auto-active SNC1 or/and its essential downstream signaling component(s) into the nucleus (Palma *et al.*, 2005). Since mutations in *MOS6* only partially suppresses *snc1* autoimmunity and *mos6* single mutants show only mild defects in basal resistance (Palma *et al.*, 2005; Roth *et al.*, 2017), other α-importins may have partially overlapping functions with MOS6.

Here, we investigated the genetic requirement of the nine *Arabidopsis α-importin* paralogs for manifestation of the autoimmune phenotype of *snc1*, and we analyzed the nuclear import complex formation of the nine α-importins with the auto-active and wildtype SNC1 proteins. We show that both protein variants of SNC1 strongly interact with MOS6/IMP-α3, whereas we detected no or only very weak interactions with the other eight *Arabidopsis* α-importins when co-expressed transiently in leaves of *N. benthamina*. The preferential association of SNC1 with MOS6/IMP-α3 is consistent with reverse-genetic analyses showing that a mutation in *MOS6/IMP-α3*, but not in any other *importin-α* gene, partially suppresses the dwarf autoimmune phenotype of *snc1* plants. In addition, *mos6* but no other *importin*-*α* single mutant is compromised in basal disease resistance. This suggests that MOS6/IMP-α3 plays a major functional role in nuclear import of SNC1 and possibly of cargo proteins involved in the regulation of basal resistance in *Arabidopsis*.

## RESULTS

### *MOS6*/*IMPORTIN-α3* is selectively required for autoimmunity of *snc1*

In *Arabidopsis* genetic screens, three mutant alleles of the nuclear transport receptor *MOS6/IMPORTIN-α3* (*IMP-α3*; *AT4G02150*) were identified as partial suppressors of constitutive immunity activated in the *NLR* gene mutant *snc1* (Zhang *et al.*, 2003; Palma *et al.*, 2005). Whereas *snc1* plants are severely stunted and have curly leaves, *snc1 mos6* double mutants are of intermediate size between the wildtype Col-0 and *snc1* and have leaves that are less curly compared to *snc1* (Palma *et al.*, 2005). Since all mutant alleles of *mos6* do not completely suppress *snc1* autoimmunity (Palma *et al.*, 2005; Wirthmueller *et al.*, 2015; Roth *et al.*, 2017), we reasoned that other *α-importins* may partially compensate for loss of *MOS6* function. The *Arabidopsis* Col-0 reference genome encodes for nine importin-α isoforms of which *IMP-α1*, *−α2*, *−α3/MOS6*, *−α4*, *−α6 and −α9* are expressed ubiquitously throughout plant tissues (Hruz *et al.*, 2008; Wirthmueller *et al.*, 2013). Figure 1 shows the phylogenetic relationships and the schematic gene structures of *Arabidopsis IMP-α1* - *IMP-α9*. In order to investigate their functional relevance for manifestation of the dwarf *snc1* autoimmune phenotype, we established a collection of T-DNA insertion mutants for all nine individual *Arabidopsis importin-α* genes that were obtained from the Nottingham Arabidopsis Stock Centre (NASC; Scholl *et al.*, 2000). If available, two independent T-DNA insertion lines for each *IMP-α* gene were ordered. Homozygous mutant lines of the individual *IMP-α* genes were isolated via PCR-based genotyping and analyzed for disruption of functional transcripts via RT-PCR, using cDNA-specific primers that flank or are downstream (upstream for *imp-α9*) of the T-DNA insertions (Figure S1). For each gene, one mutant line without detectable full-length transcripts was subsequently used for further functional analyses (Figures 1 and S1). To determine whether the *imp-α* mutants can suppress the *snc1* autoimmune growth morphology, we crossed *snc1* with each of the *imp-α* mutant lines to obtain homozygous *snc1 imp-α* double mutants in the F_2_ generation. As shown in Figure 2, only a defect in *MOS6*/*IMP-α3* but not in any of the other eight *importin-α* genes partially suppresses the stunted growth of *snc1* irrespective of the photoperiod. These genetic data indicate that *IMP-α1*, *−α2*, *−α4*, *−α5*, *−α6*, *−α7*, *−α8* and *−α9* are not individually essential for establishment of the *snc1* autoimmune phenotype, whereas *IMP-α3/MOS6* plays a prominent role in *snc1*-mediated growth retardation.

**Figure 1.**
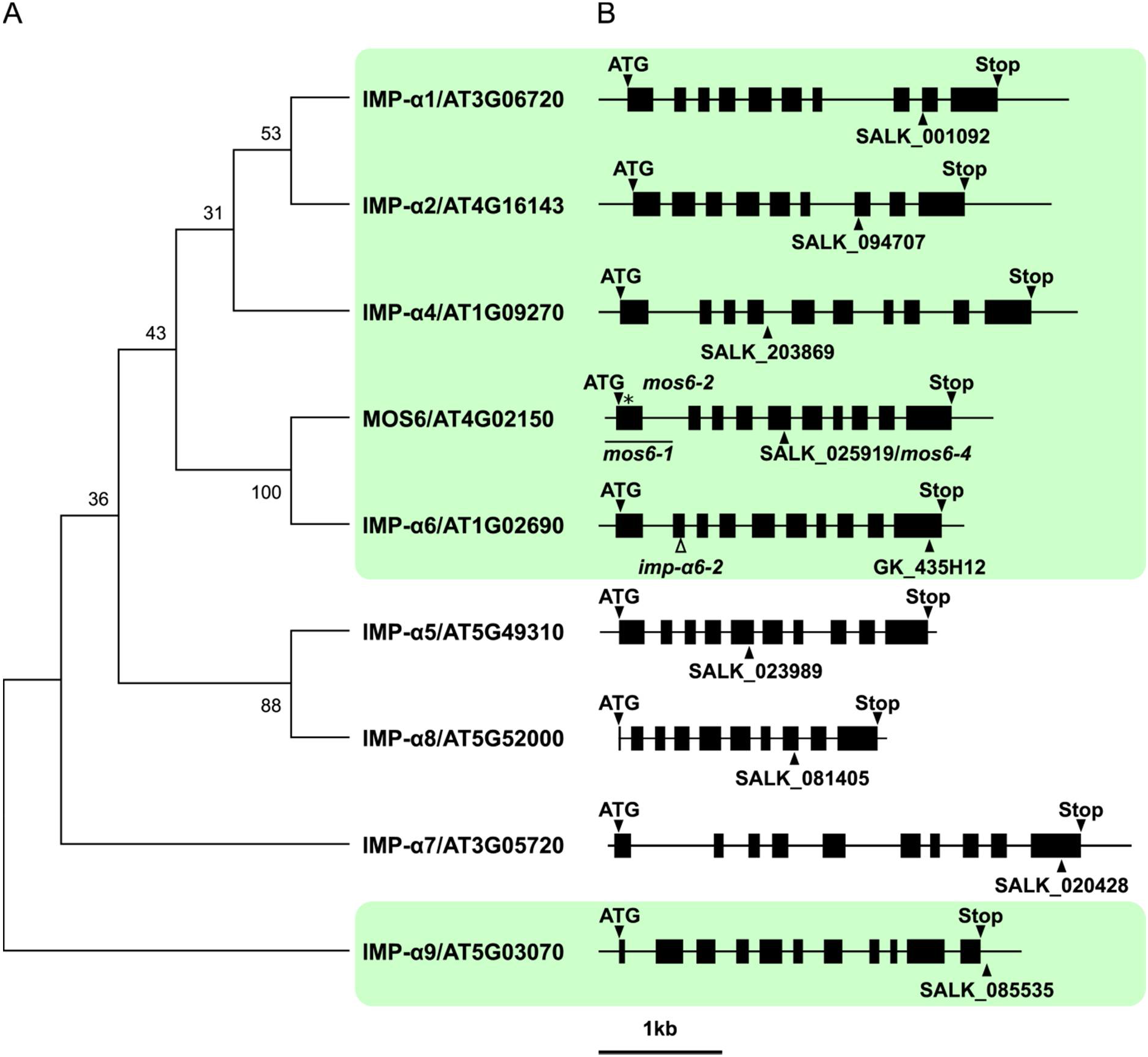
Phylogenetic tree and schematic gene structures of α-importins in *Arabidopsis*. (A) Bootstrap consensus tree using the WAG maximum likelihood method based on a manually refined MUSCLE alignment of the full length amino acid sequences performed in MEGA X (v10.0.5). IMP-α9 was used to root the tree, numbers at nodes indicate support from 100 bootstraps. α-IMPs with strong expression in rosette leaves are highlighted in green (Wirthmueller *et al.*, 2013). (B) Gene structure drawn to scale with exons as black boxes and introns as solid lines. Start (ATG) and Stop codons are indicated as triangles above, positions of the respective T-DNA insertions as triangles below gene structures. The solid line below the *MOS6* gene structure marks the approximate region of the genomic rearrangement in the *mos6-1* mutant, the deletion in the *mos6-2* mutant is indicated by an asterisk (Palma *et al.*, 2005), and the premature Stop codon in the *imp-α6-2* CRISPR/Cas9 mutant is indicated by an open triangle.

**Figure 2.**
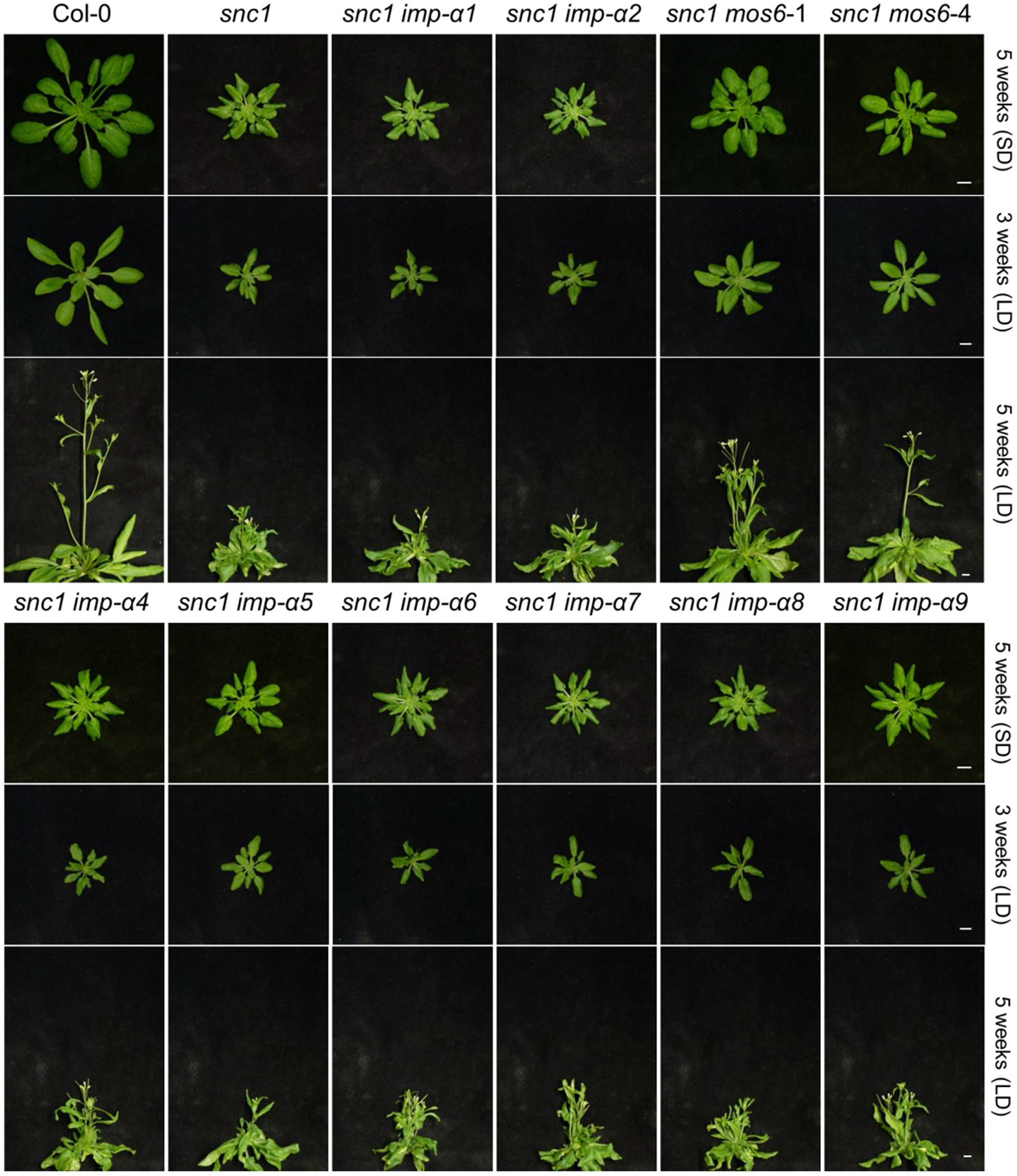
Only a loss of *MOS6* partially suppresses the *snc1*-associated stunted growth morphology. Representative images of plants grown in parallel for three or five weeks under short day (SD) and long day (LD) conditions, respectively. Scale bar = 1 cm.

*MOS6/IMP-α3* is most closely related to *IMP-α6* (Figure 1). However, the available *imp-α6* mutant allele (GK_435H12; Figure 1) that we crossed with *snc1* is located in the last exon of the gene and we therefore cannot fully exclude the presence of a partially functional IMP-α6 protein in this *imp-α6* mutant. To further investigate whether MOS6 functions redundantly with IMP-α6 in autoimmunity of *snc1*, we used CRISPR/Cas9-based genome editing to generate an *imp-α6* mutation in the *snc1 mos6-1* double mutant background. This mutation, named *imp-α6-2*, introduces a premature stop codon in the second exon of *IMP-α6/At1G02690* (Figure S2A and B). As the *imp-α6-2* mutation did not further suppress the *snc1*-associated stunted growth morphology of the *snc1 mos6-1* double mutant (i.e. *snc1 mos6-1* and *snc1 mos6-1 imp-α6-2* as well as *snc1 mos6-1 imp-α6-1* (GK_435H12) plants are of similar size and have leaves that are less curly compared to *snc1* and *snc1 imp-α6* plants; Figures 2 and S2C), our data suggest that *IMP-α6* does not function redundantly with *MOS6/IMP-α3* in the *snc1* autoimmune pathway.

### *MOS6*/*IMPORTIN-α3* is selectively required for basal resistance

Mutations in *MOS6* not only suppress the stunted growth and constitutive immunity of *snc1* (Figure 2; Palma *et al.*, 2005), but also result in enhanced disease susceptibility to the oomycete pathogen *Hyaloperonospora arabidopsidis* (*Hpa*) Noco2 and to the mildly virulent bacterial pathogen *P. syringae* pv. *tomato* (*Pst*) DC3000 lacking the effectors AvrPto and AvrPtoB (ΔAvrPto/AvrPtoB; Palma *et al.*, 2005; Wirthmueller *et al.*, 2015; Roth *et al.*, 2017). To test the contribution of the other *importin-*α genes for basal disease resistance, we inoculated the *importin-α* single mutants with *Pst* DC3000 ΔAvrPto/AvrPtoB and analyzed their disease susceptibility. For the infection assay, the Col *eds1-2* mutant was used as hyper-susceptible control, whereas the *snc1* mutant served as a control for enhanced disease resistance due to its constitutive defense activation (Zhang *et al.*, 2003; Bartsch *et al.*, 2006; Roth *et al.*, 2017). Basal resistance against *Pst* DC3000 ΔAvrPto/AvrPtoB was significantly compromised in *mos6* mutant plants when compared to the wildtype Col-0 (Figure 3). Consistent with previous data, the enhanced susceptibility of *mos6* mutant alleles was less pronounced as the complete breakdown of resistance in Col *eds1-2* plants (Wirthmueller *et al.*, 2015; Roth *et al.*, 2017). Single mutants of the other *importin-α* genes did not show significantly altered susceptibility upon inoculation with *Pst* DC3000 ΔAvrPto/AvrPtoB and were as resistant as wildtype plants (Figure 3). This suggests that among the nine *α-importins* in *Arabidopsis*, *MOS6/IMP-α3* does not only play a predominant role in *snc1*-mediated autoimmunity (Figure 2), but also in basal disease resistance (Figure 3). Accordingly, the individual disruption of *MOS6* or of any other *IMP-α* gene function is not compensated for by increased expression of the remaining functional *IMP-α* genes (Figure S3). We also did not detect an altered expression of any *IMP-α* in the *snc1* auto-immune mutant. *Vice versa*, the expression of *SNC1* was also not obviously changed in any of the *imp-α* single mutants compared to the wildtype control (Figure S3). Similarly, the *IMP*-*α* gene expression levels are not considerably altered after infection with pathogens in any of the datasets available via the Genevestigator (Hruz *et al.*, 2008) or the Bar Expression databases (Toufighi *et al.*, 2005). Together, these data strongly suggest a preferential involvement of MOS6/IMP-α3 in plant immune responses (Figures 2, 3 and S2).

**Figure 3.**
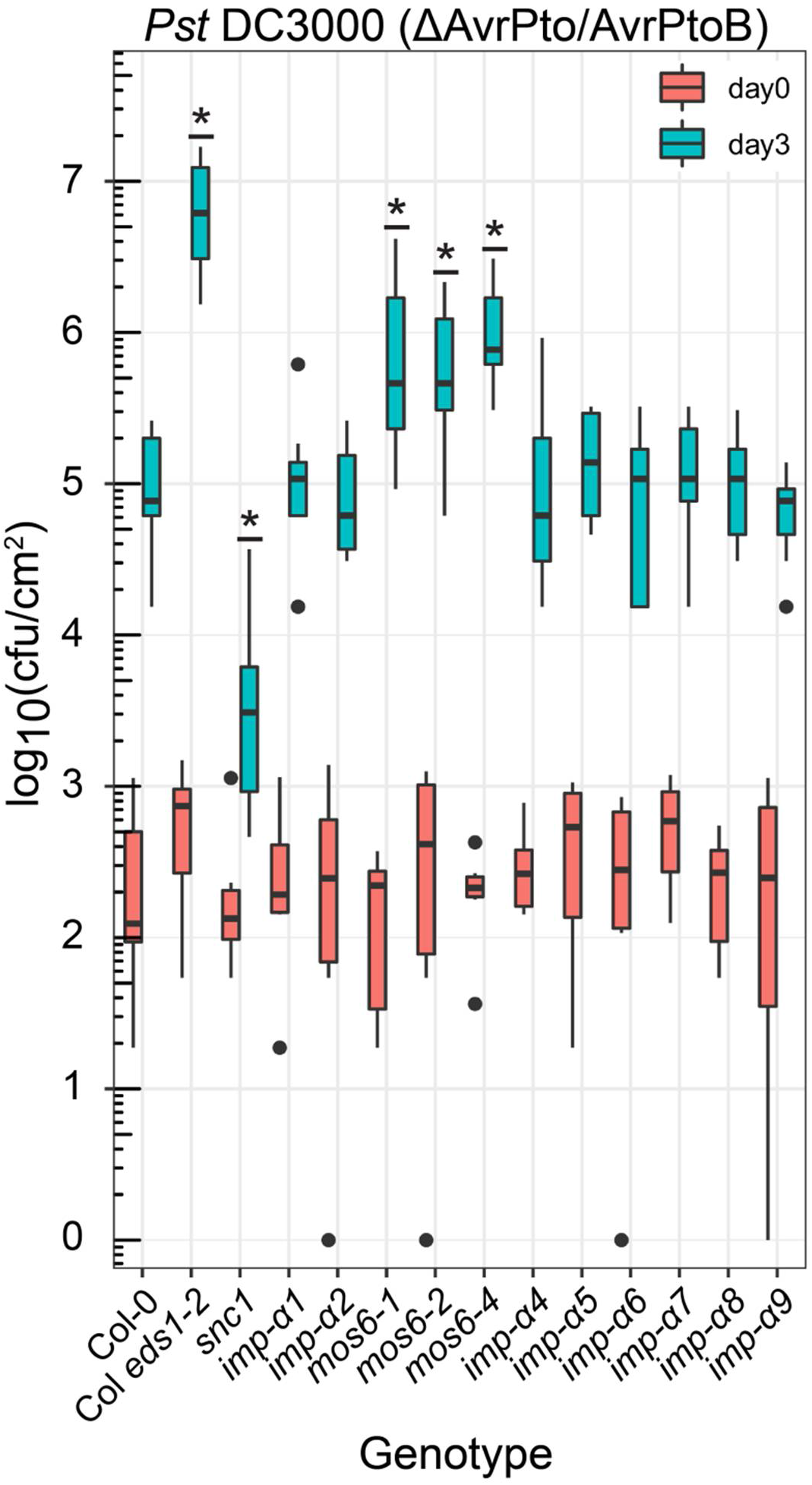
Only *mos6* alleles but no other *imp-α* mutant lines show enhanced susceptibility to mildly virulent *P. syringae*. Four week old plants of the indicated genotypes were vacuum infiltrated with a *Pst* DC3000 (ΔAvrPto/AvrPtoB) suspension of 1 × 10^5^ cfu ml^−1^. Colony-forming units (cfu) within the infiltrated plant tissues were quantified immediately (day 0), or three days after infiltration (day 3). Data is presented as boxplots (day 0: n=6; day 3: n=9), outliers are indicated as black dots, underlined asterisks indicate statistically significant differences to Col-0 (one-way ANOVA; Tukey’s test, *P* < 0.05). The experiment was repeated three times with similar results.

### Morphological characterization of *imp-α* single and higher order mutants

The overall rosette size and growth morphology of *mos6* single mutants is indistinguishable from Col-0 wildtype plants (Palma *et al.*, 2005; Roth *et al.*, 2017), suggesting that loss of *MOS6* function does not affect regular plant growth and development. To test whether this also holds true for the other *IMP-α* genes, we investigated the growth phenotypes of our *imp-α* single mutant collection. When cultivated under both short-day or long-day growth conditions, all examined *imp-α* single mutants showed a wildtype-like growth phenotype in terms of rosette/plant size and morphology, and the timing of floral transition and bolting. Only *imp-α1* plants had a marginally reduced rosette size (Figure 4). Together, this indicates that the functional loss of individual *importin-α* isoforms does not have a major impact on regular plant development.

**Figure 4.**
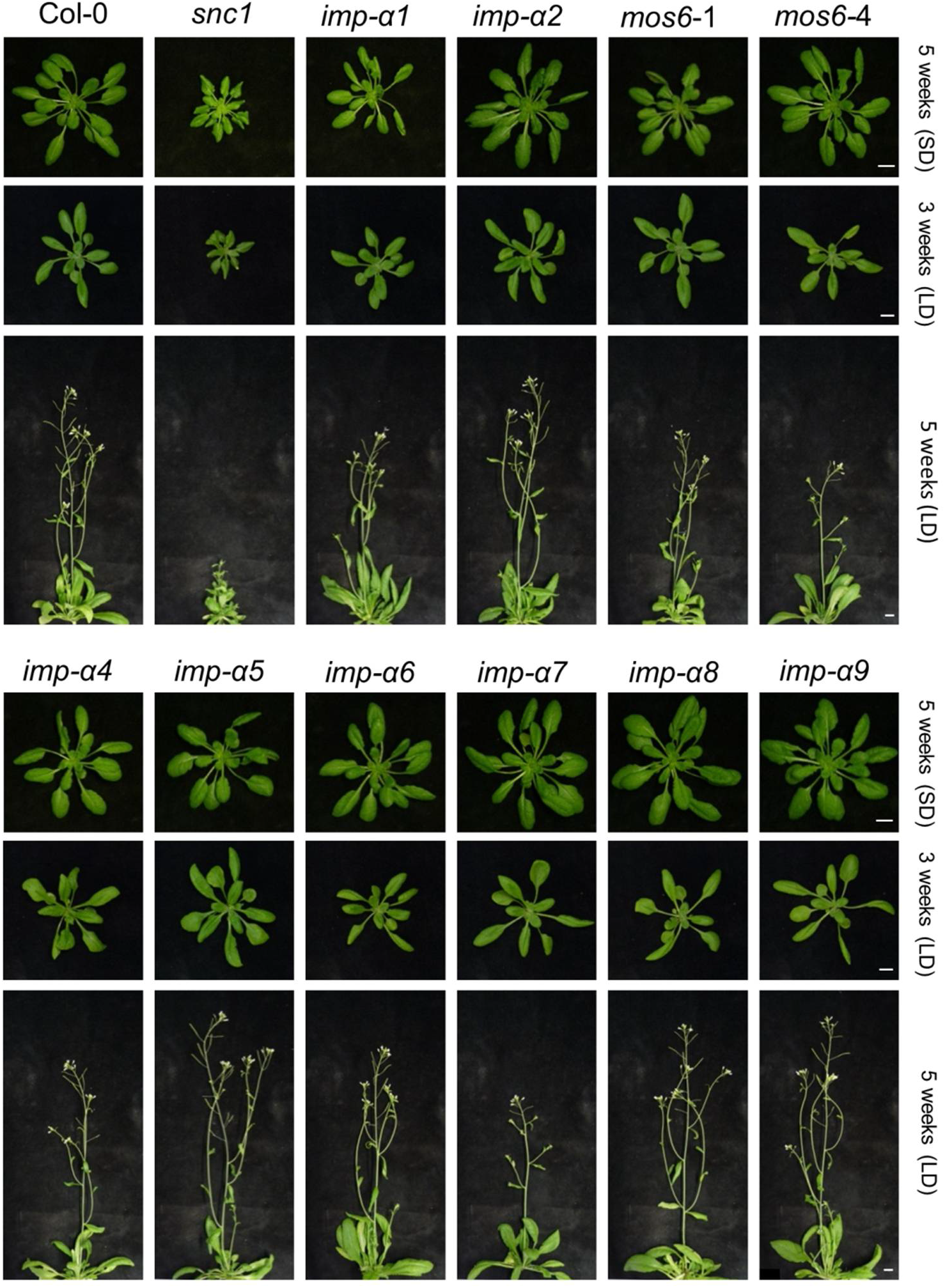
*Imp-α* single mutants do not show obvious growth defects. Representative images of plants grown in parallel for three or five weeks under short day (SD) and long day (LD) conditions, respectively. Scale bar = 1 cm.

To test whether *IMP*-*α* family members have overlapping functions in plant development, we generated several *imp-α* double and triple mutant combinations and characterized their growth phenotypes. Generally, growth of the *imp-α* double mutants was indistinguishable from Col-0 wildtype plants, except for double mutants containing the *imp-α1* allele that were smaller than wildtype plants (Figure 5). This growth reduction was most pronounced in the double mutant of the closely related *imp-α1* and *imp-α2* (Figure 1). For the *mos6 imp-α1* combinations this phenotype was most clearly seen for full-grown plants after five weeks growth under long-day conditions rather than for rosette sizes. Beside this, no other obvious morphological defects were observed for any of the *imp-α* double mutants that we investigated (Figures 5 and S2). The growth retardation that we observed for the *imp-α1 imp-α2* was even more extreme for *imp-α1 imp-α2 mos6-4* triple mutant plants (Figure 6). In particular, when grown under short-day conditions these plants were even smaller as compared to the severely stunted *snc1* control (Figure 6). The phenotypes of all other triple mutant combinations were similar to the wildtype control (Figure 6) and we did not detect an altered expression of the remaining functional *IMP-α* genes in any of the triple mutants as compared to the wildtype control (Figure S4). Together, these mutant analyses show that *IMP-α1*, *IMP-α2* and *MOS6/IMP-α3* have partially redundant functions important for regular plant growth and development. However, when we infected triple mutants containing *mos6-4* with *Pst* DC3000 ΔAvrPto/AvrPtoB, their susceptibility was similar to that of the *mos6-1* single mutant, suggesting that MOS6 plays a major functional role in maintaining the basal resistance layer to *Pst* DC3000 ΔAvrPto/AvrPtoB (Figure S5).

**Figure 5.**
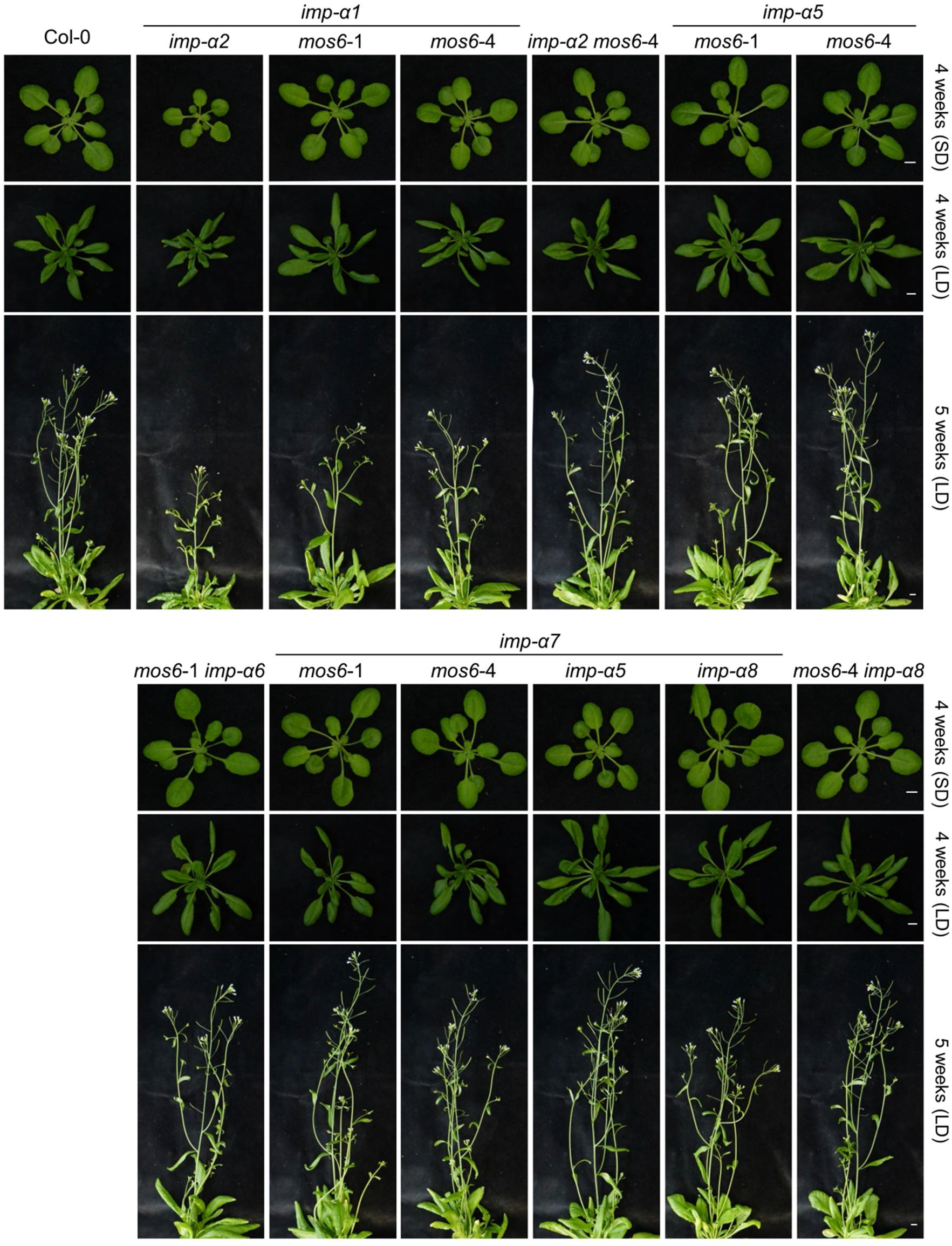
Growth morphology of *imp-α* double mutant combinations. Representative images of plants grown in parallel for four or five weeks under short day (SD) and long day (LD) conditions, respectively. Scale bar = 1 cm.

**Figure 6.**
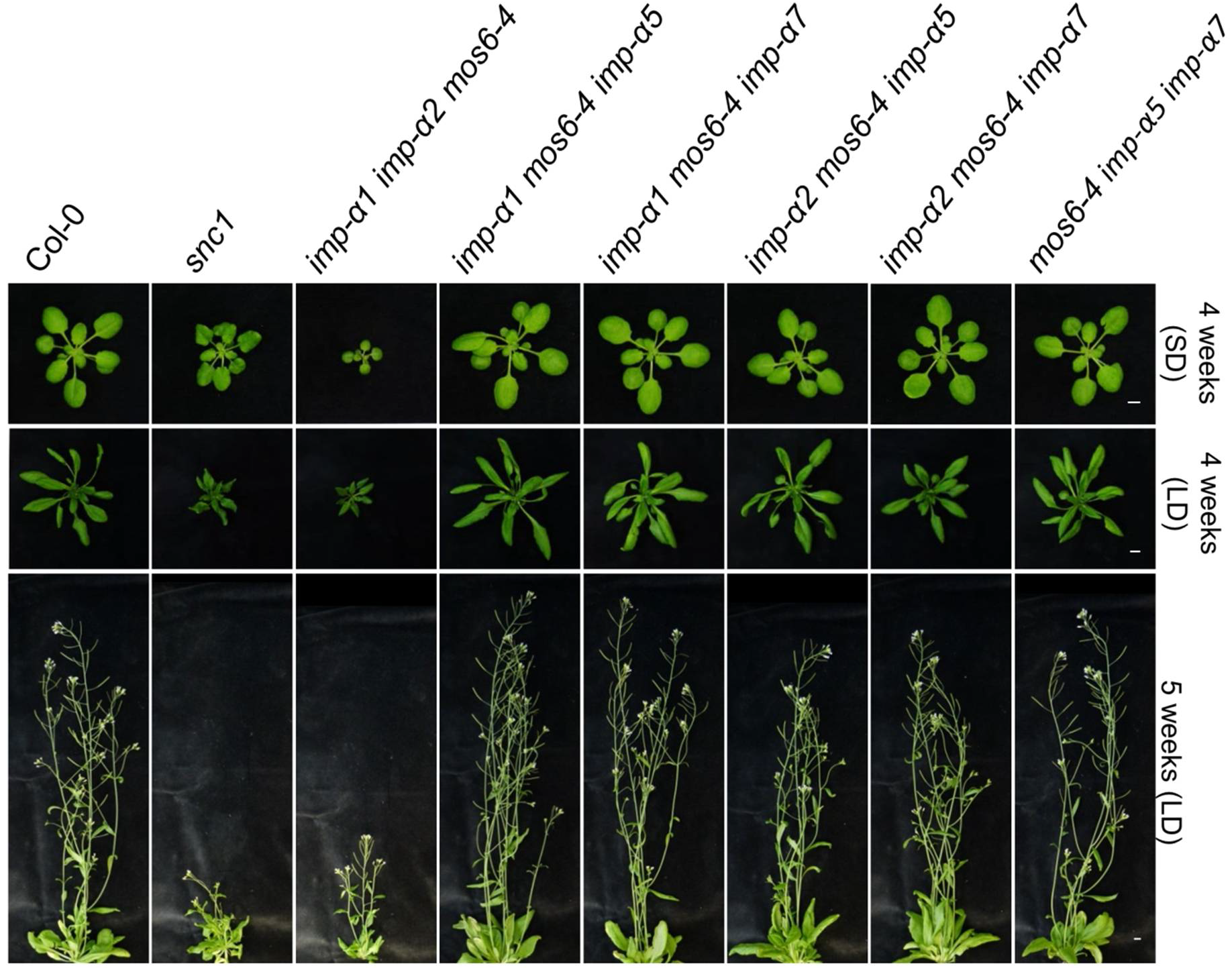
Growth morphology of *imp-α* triple mutant combinations. Representative images of plants grown in parallel for four or five weeks under short day (SD) and long day (LD) conditions, respectively. Scale bar = 1 cm.

### NLR protein SNC1 preferentially associates with MOS6/IMPORTIN-α3

The preferential genetic requirement of *MOS6/IMP-α3* for basal resistance (Figure 3) and constitutive immunity caused by the E_552_K mutation in the TNL protein SNC1 (Figure 2) is intriguing and the latter suggests that MOS6/IMP-α3 is responsible for transport of the auto-active SNC1 (SNC1^E552K^) or/and its essential downstream signaling component(s) into the nucleus.

Bioinformatic analyses with default stringency settings revealed no strong candidate sequences for an NLS in the SNC1 protein sequence (Nguyen Ba *et al.*, 2009; Kosugi *et al.*, 2009). Lowering the thresholds for NLS prediction in NLStradamus and NLSmapper identified three lower-scoring sequences that might function as NLS and mediate active nuclear import of this ~163 kDa nucleocytoplasmic TNL protein (Figure 7A and Table S1; Zhang *et al.*, 2003; Cheng *et al.*, 2009). As there are nine importin-α isoforms in *Arabidopsis* (Figure 1; Wirthmueller *et al.*, 2013), and mutations in *MOS6* suppress autoimmunity of *snc1* only partially (Figure 2; Palma *et al.*, 2005; Wirthmueller *et al.*, 2015; Roth *et al.*, 2017), other α-importins may cooperate with MOS6 in transporting auto-active SNC1^E552K^ or its downstream signal transducers. To investigate the potential interactions of MOS6/IMP-α3 and the other eight α-importins with SNC1^E552K^ *in planta*, we used *Agrobacterium*-mediated transient co-expression in leaves of *Nicotiana benthamiana eds1a-1* plants (Ordon *et al.*, 2017) and conducted co-immunoprecipitation (co-IP) analyses two days after *Agrobacterium* infiltration. Since a C-terminal 3xHA-StrepII (3xHA-SII) epitope tag of MOS6/IMP-α3 and a C-terminal fluorescent protein tag of SNC1 do not interfere with the respective protein functions (Cheng *et al.*, 2009; Roth *et al.*, 2017), we co-expressed the 3xHA-SII-tagged α-importins with mYFP-tagged SNC1^E552K^. We used the *N. benthamiana eds1a-1* mutant to investigate IMP-α/SNC1 transport complex formations, because the transient expression of both the auto-active and the wildtype SNC1 (SNC1^wt^, see below) triggers a HR-like cell death response in *N. benthamiana* wildtype plants that depends on the essential TNL downstream signaling component EDS1 (Figure S6; Wiermer *et al.*, 2005; Ordon *et al.*, 2017). Transient expression of the mYFP-tagged SNC1^E552K^ and SNC1^wt^ confirmed a previously reported nucleocytoplasmic localization (Cheng *et al.*, 2009). As expected for α-importins, we also observed a nucleocytoplasmic distribution when we transiently expressed IMP-α1 to IMP-α9 in *N. benthamina eds1a-1,* albeit IMP-α1, IMP-α2, IMP-α3/MOS6 showed a predominantly nuclear localization as reported previously for other α-importins (Figure S7 and S8; Kanneganti *et al.*, 2007; Chen *et al.*, 2018).

**Figure 7.**
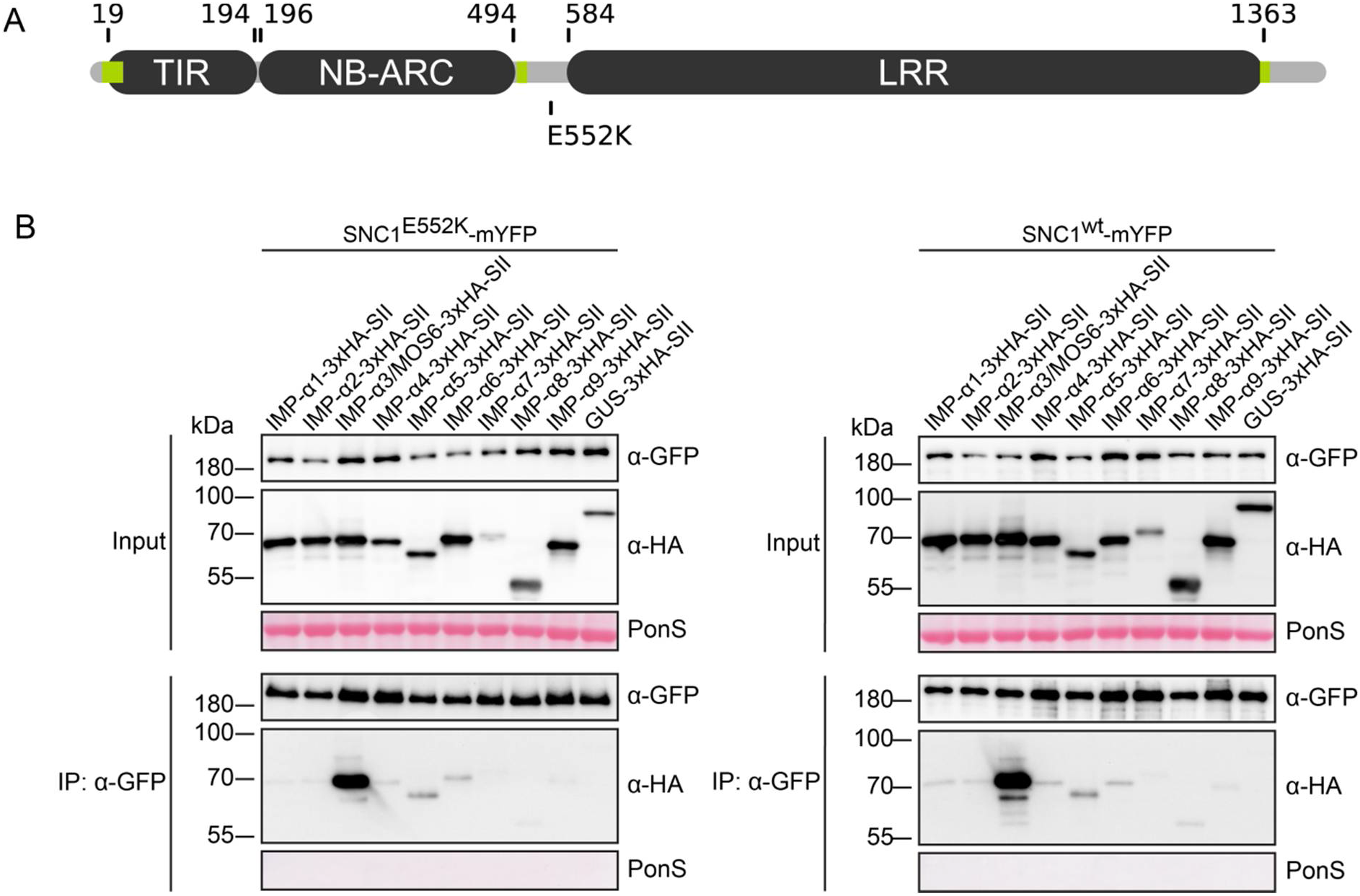
SNC1^E552K^ and SNC1^wt^ predominantly interact with IMP-α3/MOS6. (A) Schematic protein domain structure with predicted beginning and end of the respective domains, indicated by the amino acid positions within the sequence of SNC1. The auto-activating E_552_K point mutation and the predicted NLS sequences (Table S1, highlighted in green) are indicated. (B) 3xHA-StrepII (3xHA-SII)-tagged α-IMPORTINS or GUS control were transiently co-expressed in *Nicotiana benthamiana eds1a-1* with mYFP-tagged SNC1^E552K^ (left) or SNC1^wt^ (right), together with the silencing suppressor p19. Two days post-infiltration of *Agrobacterium tumefaciens*, mYFP-tagged proteins were immunoprecipitated using GFP-Trap^®^ magnetic agarose beads (IP: α-GFP). Co-immunoprecipitation of 3xHA-SII-tagged IMP-α proteins was detected by α-HA immunoblots. The upper two blots show total protein extracts (Input) probed with α-GFP and α-HA, respectively. Ponceau S (PonS) staining of the membrane was used to monitor loading. Similar results were obtained in three independent experiments.

Using GFP-Trap^®^ magnetic agarose beads (Chromotek) for immunoprecipitation of SNC1^E552K^-mYFP and subsequent detection of co-purifying 3xHA-SII-tagged α-importins by anti-HA western blot analysis revealed that the auto-active SNC1 strongly interacts with MOS6/IMP-α3, whereas we detected only very weak interactions with the other α-importins (Figure 7B, left). As the E_552_K mutation of the auto-active SNC1 protein variant is located in close proximity to one of the predicted NLSs of SNC1 (Figure 7A), we investigated whether the mutation might influence recognition specificities by importin-α proteins. Therefore, we also analyzed the interaction of the wildtype SNC1 protein (SNC1^wt^) with the nine *Arabidopsis* α-importins. As shown in Figure 7B, the association pattern of the wildtype SNC1 protein was highly similar to the auto-active protein variant, i.e. SNC1^wt^ strongly interacts with MOS6/IMP-α3, but only very weakly associates with the other importin-α isoforms. These analyses show that both the auto-active and wildtype SNC1 proteins preferentially interact with MOS6/IMP-α3 *in planta* and suggest that, among the nine α-importins in *Arabidopsis*, MOS6 is the main isoform that mediates the import of SNC1 into the nucleus. The considerably weaker associations between SNC1 and the other isoforms may collectively provide sufficient nuclear import of SNC1^E552K^ to partially compensate for loss of MOS6 function in *snc1*-mediated immunity.

## DISCUSSION

Nuclear translocation and accumulation of certain NLR proteins, signal transducers and/or transcription factors are important regulatory steps in controlling diverse cellular defense pathways in plants (Gu, 2018). Cytoplasmic proteins containing cNLSs are recruited to the nuclear transport machinery via importin-α transport adapters that recognize and bind the exposed cNLSs of their cargos. The sequenced genomes of higher plants usually encode several importin-α isoforms, suggesting either that subsets of α-importins function redundantly or that multiple isoforms have evolved to confer preferential nuclear entry of specific cargos (Wirthmueller *et al.*, 2013). The latter functional diversification may arise from tissue-specific and/or temporally distinct expression patterns, or from isoform-specific affinities for particular cargo substrates.

In *Arabidopsis*, autoimmunity of the *NLR* gene mutant *snc1* partially depends on *MOS6*/*IMP-α*3, one of nine members of the *importin-α* gene family (Palma *et al.*, 2005; Wirthmueller *et al.*, 2013). Here, we made use of the dwarf morphology of *snc1* to investigate the individual contribution of the other eight *importin-α* genes for manifestation of this characteristic growth phenotype. We show that *MOS6* plays an eminent role in *snc1*-mediated growth retardation, since the other *importin-α* genes are not individually essential for establishment of this autoimmune phenotype (Figure 2). However, mutations in *MOS6* do not fully suppress the dwarfism of *snc1* (Figure 2; Palma *et al.*, 2005). We therefore cannot exclude the possibility that one or several other *α-importin(s)* may partially compensate for loss of *MOS6* function in *snc1*-dependent autoimmunity, albeit the higher order mutant combinations of *mos6* that we investigated do not further impair basal defense responses, and the dwarfism of *snc1* is not further suppressed in *snc1 mos6 imp-α6* triple mutants as compared to *snc1 mos6* double mutant plants (Figure S2). Alternatively, an importin-α independent nuclear transport pathway may operate redundantly with MOS6. For instance, some NTRs of the importin-β family are capable to directly bind to cargo proteins and mediate nuclear import independently of importin-α (Christie *et al.*, 2016; Zhang *et al.*, 2017; Liu *et al.*, 2019). Given that the SNC1 sequence does not harbor strong candidates for a cNLS, it remains possible that direct binding to MOS6 is mediated by an atypical NLS. It is also conceivable that SNC1 might employ a piggy-back mechanism and binds to MOS6 indirectly via an NLS-containing interaction partner or specific adapter protein, as shown for nuclear import of *Arabidopsis* phyA (Genoud *et al.*, 2008).

Our co-immunoprecipitation assays suggest that, among the nine importin-α family members in *Arabidopsis*, MOS6 is the main nuclear import receptor of SNC1, providing evidence for functional specialization of MOS6 (Figure 7). Support of this idea comes from a report showing that nuclear accumulation of GFP-tagged SNC1-4 (a variant of the auto-active SNC1^E552K^ that harbors an additional E_640_K mutation) is affected in *mos6* mutant protoplasts when compared to protoplasts of *Arabidopsis* wildtype plants (Zhu *et al.*, 2010b). To obtain insights into possible specificity determinants that could explain the high preference of SNC1 for MOS6 as NTR, we analyzed the crystal structure of the MOS6 ARM repeat domain (Wirthmueller *et al.*, 2015) for amino acid polymorphisms that are not shared with any other α-importin isoform. At the MOS6 major and minor NLS binding sites, all core residues that can be predicted to make direct contact to cNLS are conserved and therefore cannot explain specificity for SNC1 (Wirthmueller *et al.*, 2013; 2015). The only two surface-exposed residues within the MOS6 inner solenoid that are not shared with any other *Arabidopsis* α-importin are N_275_ and V_444_. Based on crystal structures of rice importin-α in complex with cNLS peptides (Chang *et al.*, 2012; 2013), neither of these two residues directly contributes to the NLS binding sites. However, N_275_ maps to the third helix of MOS6 armadillo repeat 5 and is located in proximity to the cleft that accommodates the K in the P4’ position of cNLS peptides binding to the minor NLS binding site (Marfori *et al.*, 2011). Whether these polymorphic residues outside of the core NLS binding grooves can explain the predominant role of MOS6 as NTR for SNC1 is currently not known. However, from our protein interaction assays we conclude that the considerably reduced associations of SNC1 with the other importin-α isoforms may collectively be sufficient for transporting enough of the SNC1^E552K^ pool into the nucleus to partially compensate for loss of MOS6 function in *snc1 mos6* plants (Figures 2 and 7). This would explain why *mos6* only partially suppresses *snc1*, and why the additional mutation of the closely related *IMP-α6* in *snc1 mos6 imp-α6* triple mutants is not sufficient to further suppress the *snc1*-associated growth retardation of *snc1 mos6* double mutants (Figure S2C).

It should be noted that our conclusion is based on the formation of SNC1/IMP-α transport complexes in the heterologous expression system *N. benthamiana*. We therefore cannot fully exclude that the differential complex formations are, at least in part, due to competition by other NLS-cargos in the cytosol of *N. benthamiana* cells, yet our reverse-genetic analysis in *Arabidopsis* are consistent with the preferential co-purification of MOS6 with immunoprecipitated SNC1 (Figure 7). Our co-IP results further suggest that the E_552_K mutation located in proximity to one of the predicted NLSs of SNC1 does not obviously alter the importin-α binding affinities/specificities for the auto-active SNC1^E552K^ as compared to SNC1^wt^ (Figure 7). Nuclear transport rates of cargos are directly related to their binding affinities for their import receptors (Hodel *et al.*, 2006; Timney *et al.*, 2006; Christie *et al.*, 2016). Therefore, it is unlikely that the *snc1* autoimmune phenotype induced by the E_552_K mutation is simply caused by altered nuclear import rates of the SNC1^E552K^ protein.

Considering near identity in the core residues forming the NLS-binding sites of MOS6 and the leaf-expressed isoforms α1, α2, α4 and α6 (Wirthmueller *et al.*, 2015), it is intriguing that MOS6 preferentially associates with SNC1 (Figure 7), and that the knock-out of a single *importin-α* gene causes a specific phenotype such as the immunity defects of *mos6* (Figures 2, 3, S2 and S5). In another example, Bhattacharjee *et al.* (2008) reported that the loss of *IMP-α4* but not of other *importin-α* genes impairs host transformation by *Agrobacterium tumefaciens*, albeit several importin-α isoforms are able to interact with the NLS-containing *A. tumefaciens* effectors VirD2 and VirE2 that mediate nuclear translocation of the T-complex. *IMP-α4* has the highest expression level in *Arabidopsis* roots compared to the other *α-importin* genes (Wirthmueller *et al.*, 2013). Significantly, Bhattacharjee *et al.*, (2008) also show that the *imp-α4* phenotype can be complemented by overexpression of not only *IMP-α4*, but also of several other isoforms. This suggests that although IMP-α4 appears to be the most crucial isoform for transfer of the T-complex, other isoforms can compensate the loss of IMP-α4 function when their cellular abundance is increased (Bhattacharjee *et al.*, 2008). Thus, nuclear import kinetics are influenced not only by the affinity of a particular cargo for the NTR, but can also be modified by the cytoplasmic concentrations of both the NTRs and the cargo proteins (Timney *et al.*, 2006; Bhattacharjee *et al.*, 2008; Cardarelli *et al.*, 2009; Wirthmueller *et al.*, 2015).

In plants, there are more prior examples of importin-α cargo selectivity as well as of redundancy (Jiang *et al.*, 2001; Kanneganti *et al.*, 2007; Bai *et al.*, 2008; Wirthmueller *et al.*, 2015; Gerth *et al.*, 2017; Roth *et al.*, 2017; Chen *et al.*, 2018; Contreras *et al.*, 2019). Accordingly, in addition to the selective role of MOS6/IMP-α3 in plant (auto-)immunity (Figures 2, 3, 7, S2 and S5), our genetic analysis also revealed that *MOS6* has partially overlapping functions with *IMP-α1* and *IMP-α2* in regular plant growth and development (Figure 6). The molecular basis for this functional specialization/redundancy in nuclear import pathways remains elusive. We speculate that polymorphic residues outside of the core NLS binding grooves of MOS6/IMP-α3 may explain the predominant role as NTR for SNC1 and possibly other cargos involved basal defense regulation.

Nuclear import rates are regulated at several levels, including post-translational modifications that can modulate cargo binding affinities by introducing direct changes to the NLS or by blocking importin-α/cargo interactions through intra- or intermolecular masking of NLSs (Christie *et al.*, 2016). For example, phosphorylation of residues in vicinity of NLSs can result in cytoplasmic retention of the modified cargo variant by preventing its association with importin-α (Rona *et al.*, 2013; Helizon *et al.*, 2018). *Vice versa*, phosphorylation of residues in the linker region of bipartite NLSs can enhance the affinity of an NLS for importin-α binding and increase nuclear translocation rates (Hübner *et al.*, 1999; Christie *et al.*, 2016). Whether the SNC1/IMP-α affinities and/or binding specificities in *Arabidopsis* are modulated via post-translational modifications according to the actual cellular needs in response to environmental stimuli remains to be determined.

## EXPERIMENTAL PROCEDURES

### Plant material and growth conditions

*Arabidopsis thaliana* plants were grown on soil in environmentally controlled chambers with 65 % relative humidity. Short day conditions (SD) with an 8/16 h light/dark regime or long day conditions (LD) with a 16/8 h light/dark regime at 22/18 °C were used for growth morphology assessments. *Nicotiana benthamiana* plants were grown under a 16/8 h light/dark regime at 25/22 °C, 4-5 week old plants were used for transient expression assays. The *N. benthamiana eds1a-1* mutant was previously described (Ordon *et al.*, 2017). *A. thaliana* T-DNA insertion lines were obtained from the Nottingham Arabidopsis Stock Centre (NASC, http://arabidopsis.info). PCR-based genotyping using T-DNA flanking primers were used to identify and isolate homozygous mutants (Table S2). The *snc1* (Li *et al.*, 2001), Col *eds1*-2 (Bartsch *et al.*, 2006), *mos6-1* and *mos6*-2 (Palma *et al.*, 2005), and *mos6-4* (Wirthmueller *et al.*, 2015) mutants were previously described. For the generation of *imp-α6-2* mutant lines, transgenic plants were generated by transforming *snc1 mos6-1* double mutant with *Agrobacterium tumefaciens* strain GV3101 pMP90RK carrying a binary vector for CRISPR/Cas9-based gene editing (see below) by floral-dip (Clough and Bent, 1998). Homozygous *snc1 mos6-1 imp-α6-2 mutants* were isolated using a T7-endonuclease assay before determining the mutation via sequencing of the PCR amplified target region. *mos6-1 imp-α6-2* double and *imp-α6-2* single mutant plants were isolated by PCR-based genotyping after backcrossing of *snc1 mos6-1 imp-α6-2* with Col-0 wildtype.

### *Pseudomonas* infection assay

Plant infection assays were performed as previously described (Roth *et al.*, 2017). Briefly, rosette leaves of 5-week-old soil grown plants were vacuum-infiltrated with a suspensions of *Pst* DC3000 ΔAvrPto/AvrPtoB (Lin and Martin, 2005) with a bacterial density of 1 × 10^5^ cfu ml^−1^ in 10 mM MgCl_2_ and 0.001 % (v/v) Silwet L-77. Titers were determined 1 h (d0) and 3 days (d3) after infiltration.

### Construction of plasmids

The Gateway compatible binary destination vectors pXCSG-mYFP or pXCSG-3xHA-StrepII were used for transient expression assays in *N. benthamiana* (Witte *et al.*, 2004; Feys *et al.*, 2005; García *et al.*, 2010). Genomic sequences were PCR amplified using gene specific primers listed in Table S2 and cloned into pENTR/D-TOPO (ThermoFisher Scientific, https://www.thermofisher.com) entry vectors. Sequenced entry vectors were used in LR-reactions with the respective destination vectors to receive expression vectors. The binary vector for CRISPR/Cas9-based gene editing is derived from pHEE401 (Wang *et al.*, 2015) and was provided by C. Thurow and C. Gatz (University of Goettingen). The sgRNA was created via golden-gate cloning of annealed complementary oligos 5’-**GATT**GAGACTTACAATTGGAGGCGA-3’ and 5’-**AAAC**TCGCCTCCAATTGTAAGTCTC-3’ determining the target site (overhangs for golden gate ligation sites are marked with bold letters). All sequence confirmed vectors were transformed into *Agrobacterium tumefaciens* strain GV3101 pMP90RK using electroporation.

### Transient expression in *N. benthamiana*

Agrobacteria were grown over night, harvested by centrifugation, resuspended in infiltration buffer (10 mM MgCl_2_, 10 mM MES pH 5.5, 150 μM acetosyringone) and incubated for 2-3 h at room temperature. For co-expression, *Agrobacterium* strains carrying the desired constructs were mixed with infiltration buffer to obtain a respective final optical density (OD_600_) of 0.3 for each strain. Infiltration into the abaxial side of 4-5 week old *N. benthamiana* leaves was conducted using a needleless syringe. The silencing suppressor p19 was co-infiltrated in all transient expression experiments. For cell death assays, the infiltrated leaf areas were photographed 5 days after infiltration. Co-immunoprecipitation and confocal laser scanning microscopy were performed 2 days after infiltration of Agrobacteria.

### Co-immunoprecipitation, total protein extracts and immunoblot analysis

For co-immunoprecipitation, infiltrated *N. benthamiana* leaf tissue was harvested, frozen in liquid nitrogen, homogenized by using a TissueLyser LT (Qiagen) and stainless steel beads and mixed with protein extraction buffer (250 mM sucrose, 100 mM HEPES-KOH, pH 7.5, 5 % (v/v) glycerol, 2 mM Na_2_MoO_4_, 25 mM NaF,10mM EDTA, 1mM DTT, 0.5 % (v/v) Triton X-100, plant protease inhibitor cocktail (#P9599, Sigma)). Cell debris was removed by centrifugation at 17.000 *g* and filtering through a 95-μm nylon mesh. 7.5 μl GFP-trap magnetic agarose beads (Chromotek) were equilibrated in protein extraction buffer, added to the total protein extract and incubated for 3 h at 4°C under constant rotation. Magnetic agarose beads were isolated using a magnetic rack and washed 3 times in 1 ml extraction buffer. Immunoprecipitated proteins were eluded by boiling in 4x SDS loading dye (250 mM Tris-HCl (pH 6.8), 8 % (w/v) SDS, 40 % (v/v) Glycerol, 0.04 % (w/v) Bromophenol blue, 400 mM DTT). Input samples were mixed with SDS loading dye before adding GFP-trap beads. For total protein extracts, homogenized plant material was boiled directly in 2x SDS loading dye, debris was removed by centrifugation at 17.000 *g* and the supernatant was used for immunoblot analysis. Total protein extracts and input or IP samples were separated on 7.5 % or 10 % SDS polyacrylamide gels. Proteins were transferred onto nitrocellulose membranes (Amersham Protran, 0.45 μm; GE Healthcare Life Sciences) and incubated with primary α-GFP (monoclonal, #11814460001, Roche) or α-HA (H9658; Sigma-Aldrich) antibody.

The secondary goat anti-mouse IgG-poly-HRP (polyclonal, #32230; ThermoFisher Scientific) antibody was incubated and detected using SuperSignal West Femto chemiluminescence substrate (#34095; ThermoFisher Scientific) on a ChemiDoc imaging system (BioRad).

### Confocal laser scanning microscopy

Microscopy was performed 2 days post infiltration of Agrobacteria with leaf discs embedded in water using a 20x/0.70 objective (PL APO, CS) of a Leica TSC-SP5 confocal laser-scanning microscope controlled by Leica LAS AF software. YFP was excited using 514 nm of an argon laser line and mCherry was excited using 561 nm of a DPSS laser. Emitted fluorescence was detected at 525 – 555 nm for YFP and 580 – 620 nm for mCherry using Leica HyD detectors. Images were sequentially scanned (512×512 at 400 Hz). Channels were merged using ImageJ (Schindelin *et al.*, 2012).

### RNA isolation and RT-PCR analyses

Total RNA was isolated as described in Genenncher *et al.* (2016). Briefly, RNA was isolated from soil grown plants using Trizol. Reverse transcription was performed using RevertAid H Minus reverse transcriptase (Fermentas) and an oligo(dT)_18_V primer at 42 °C in a 20 μL reaction volume with 1.5 μg of DNaseI-treated RNA as input. RT-PCRs to analyze disruption of full length transcripts were performed using primers listed in Table S2.

### *In silico* analyses

Protein domain predictions of SNC1 were performed using the InterProScan 5 web tool (Zdobnov and Apweiler, 2001). In addition, a Phyre2 (Kelley *et al.*, 2015) homology model was used to determine the beginning of the unstructured region after the MHD-like motif to assess the boundary of the NB-ARC domain (Steele *et al.*, 2019). NLS-predictions were performed using NLStradamus (Nguyen Ba *et al.*, 2009) and NLSmapper (Kosugi *et al.*, 2009). Exon-intron structures for schematic gene models are based on the Araport11 genome annotation (Cheng *et al.*, 2017). Phylogenetic analysis was performed using the maximum likelihood method and Whelan and Goldman model (WAG; Whelan and Goldman, 2001) with 3 Gamma categories in MEGA X (v10.0.5) (Kumar *et al.*, 2018).

### Statistical analyses

Statistical differences were determined using R (http://www.R-project.org). One-way analysis of variance (ANOVA) followed by Tukey’s HSD was performed after analysis of normal distribution using Shapiro-Wilk’s test and analysis of variance using Levene’s test on log-transformed data. Statistically significant differences (*P* < 0.05) are marked by asterisks. The data is presented using boxplots, with outliers defined as data points outside 1.5 times the interquartile range above/below the upper/lower quartile.

## ACKNOWLEDGEMENTS

We thank Xin Li (UBC Vancouver) for *mos6-1* and *mos6-2* seeds, Johannes Stuttmann (MLU Halle-Wittenberg) for *N. benthamiana eds1a-1* seeds, C. Gatz and C. Thurow (University of Goettingen) for binary CRISPR/Cas9 vectors and Jane Parker (MPIPZ Cologne) for pXCSG-3xHA-StrepII and pXCSG-mYFP destination vectors. We acknowledge Elena Petutschnig and Volker Lipka (University of Goettingen, Central Microscopy Facility of the Faculty of Biology and Psychology) for support and access to a Deutsche Forschungsgemeinschaft (DFG)-funded confocal microscopy platform (DFG grant INST 186/824-1 FUGG to V.L.). This research was funded by the DFG IRTG 2172 “PRoTECT” program of the Goettingen Graduate Center of Neurosciences, Biophysics, and Molecular Biosciences and the DFG research grants WI3208/4-2 and WI 3208/5-1 to M.W.

## AUTHOR CONTRIBUTIONS

D.L., C.R. and M.W. conceived and designed the experiments. D.L., C.R., S.A.K., J.M., D.H., J.A., B.F.H., Q.Y., S.K., M.K. and A.G. performed the experiments. D.L., C.R., L.W. and M.W. analyzed and discussed the data. D.L., L.W. and M.W. wrote the manuscript with contributions from the other authors.

## SUPPORTING INFORMATION

**Figure S1.** Schematic gene structures of *Arabidopsis* α-importins and their gene expression in wildtype and respective mutants as investigated by RT-PCR analysis.

**Figure S2.** Independent mutant alleles of *imp-α6* do not further suppress the *snc1*-associated stunted growth morphology.

**Figure S3.** *Imp-α* single mutants do not show obviously altered expression of the remaining functional *IMP-α* genes or *SNC1*.

**Figure S4.** *Imp-α* triple mutants or *snc1* mutants do not show obviously altered expression of the remaining functional *IMP-α* genes.

**Figure S5.** Only the *mos6* allele contributes to immunity against mildly virulent *Pst* in *imp-α* triple mutant lines.

**Figure S6.** SNC1^wt^ and SNC1^E552K^ can be transiently expressed to detectable levels in the *Nicotiana benthamiana eds1a-1* mutant without induction of a cell death response.

**Figure S7.** Transiently expressed α-IMPORTINS, SNC1^wt^ and SNC1^E552K^ show a nuclear-cytoplasmic localization in *Nicotiana benthamiana*.

**Figure S8.** Accumulation of full length mYFP-tagged IMPORTIN-α proteins upon transient expression in *N. benthamiana*.

**Table S1.** NLS predictions for SNC1.

**Table S2.** Primers used in this study.

