## Supplementary Figures and Tables for "Functional requirement of the *Arabidopsis* importin-α nuclear transport receptor family in autoimmunity mediated by the NLR protein SNC1"

**SUPPORTING INFORMATION**

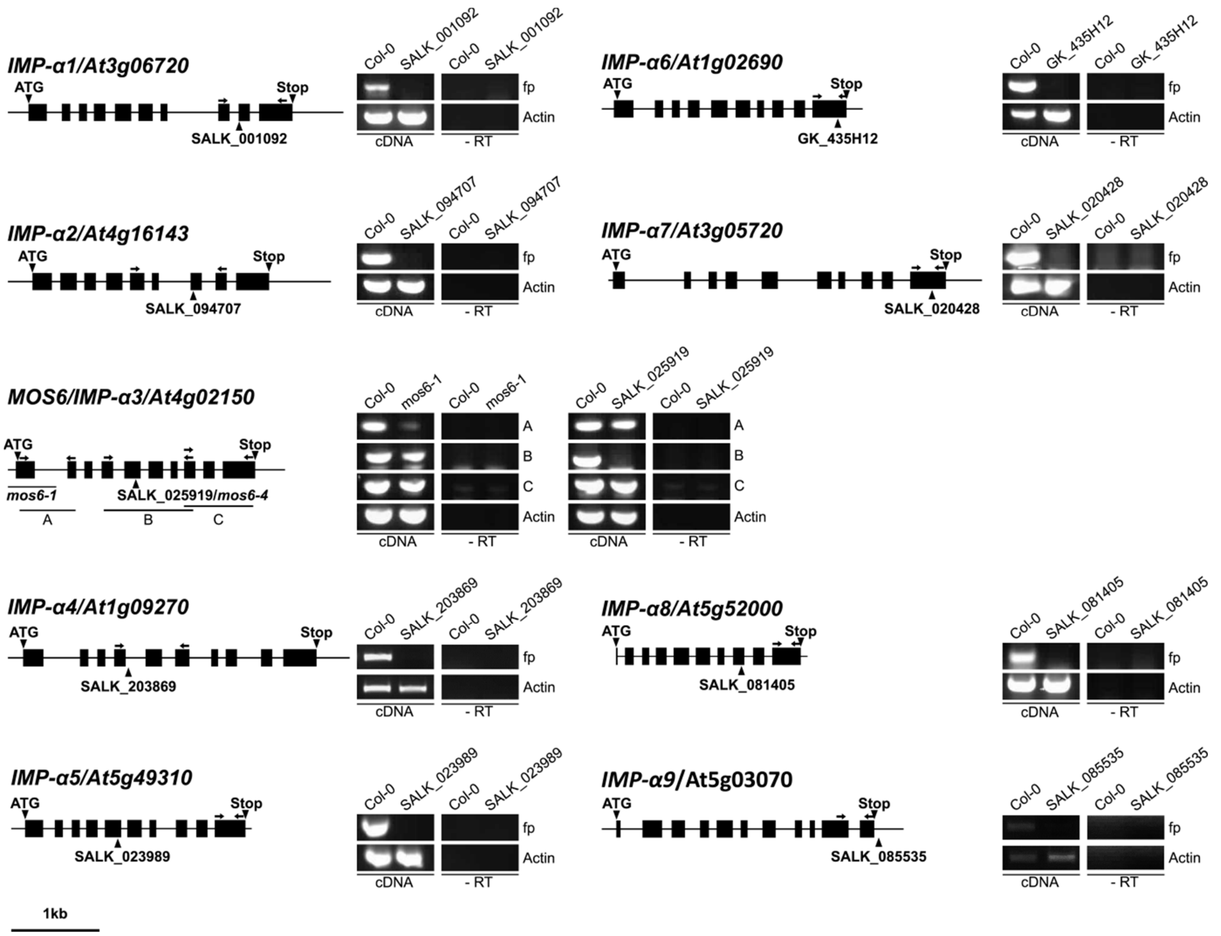


**Figure S1.** Schematic gene structures of *Arabidopsis* $\alpha$-*importins* and their gene expression in wildtype and respective mutants investigated by RT-PCRs. Gene structures and *imp-*$\alpha$ mutants used in this study (Figure 1), drawn to scale with exons as black boxes and introns as solid lines. Start (ATG) and Stop codons are indicated as triangles above, positions of the respective T-DNA insertions as triangles below gene structures. The solid line below the *MOS6* gene structure marks the approximate region of the genomic rearrangement in *mos6-1* (Palma *et al.*, 2005). Arrows above gene structure mark the position of primers used to amplify fragments from cDNA by RT-PCRs to investigate disruption of functional transcripts. For analysis of *mos6* transcripts, different primer combinations are labeled as A, B and C. -RT samples without reverse transcription were used as controls for gDNA contamination. Col-0 cDNA was used as a wild type control. PCR products were separated by agarose gel electrophoresis and stained by ethidium bromide.

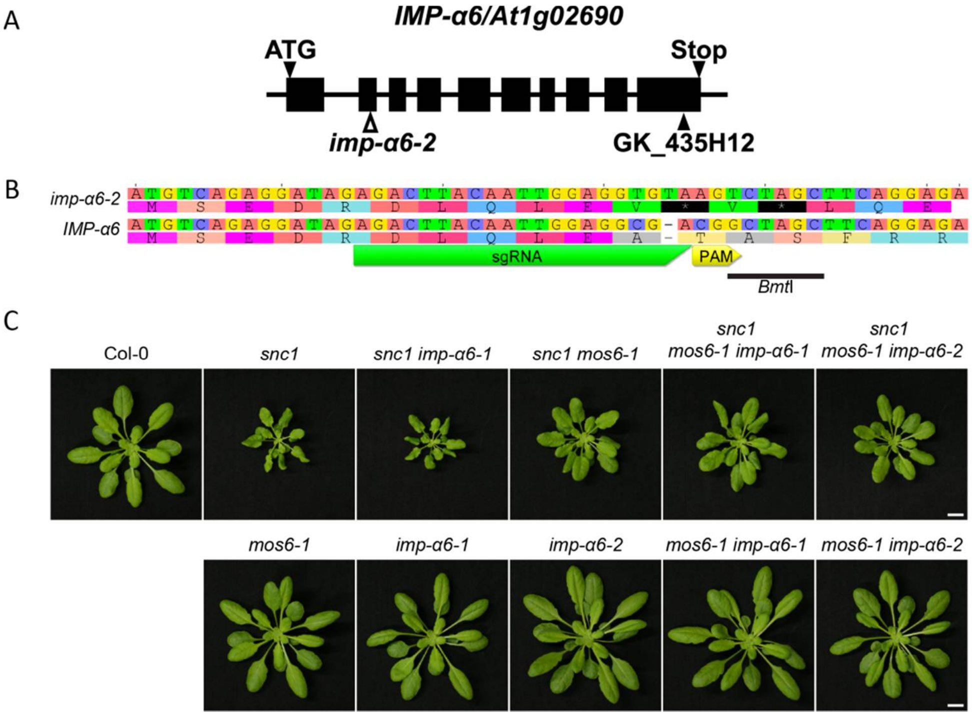


**Figure S2.** Independent mutant alleles of *imp-α6* do not further suppress the *snc1*-associated stunted growth morphology. (A) Schematic overview of the *IMP-α6*/AT1G02690 gene drawn to scale with exons as black boxes and introns as solid lines. Start (ATG) and Stop codon are indicated as black triangles above, positions of the T-DNA insertion as black triangle below gene structure. The position of the CRISPR/Cas9 target site of *imp-α6-2* is indicated by an open triangle below the gene structure. (B) Target site position of the single guide RNA (sgRNA) with protospacer adjacent motif (PAM) within the second exon of *IMP-α6*/AT1G02690. The CRISPR/Cas9 induced gene editing that introduces premature stop codons (marked by an asterisk in the translation) and abolishes a *Bmt*I restriction site that was used for PCR-based genotyping of *imp-α6-2* mutant plants, is indicated. (C) Representative images of plants of the indicated genotypes grown in parallel on soil for five weeks under short day (SD) conditions. Scale bar = 1 cm.

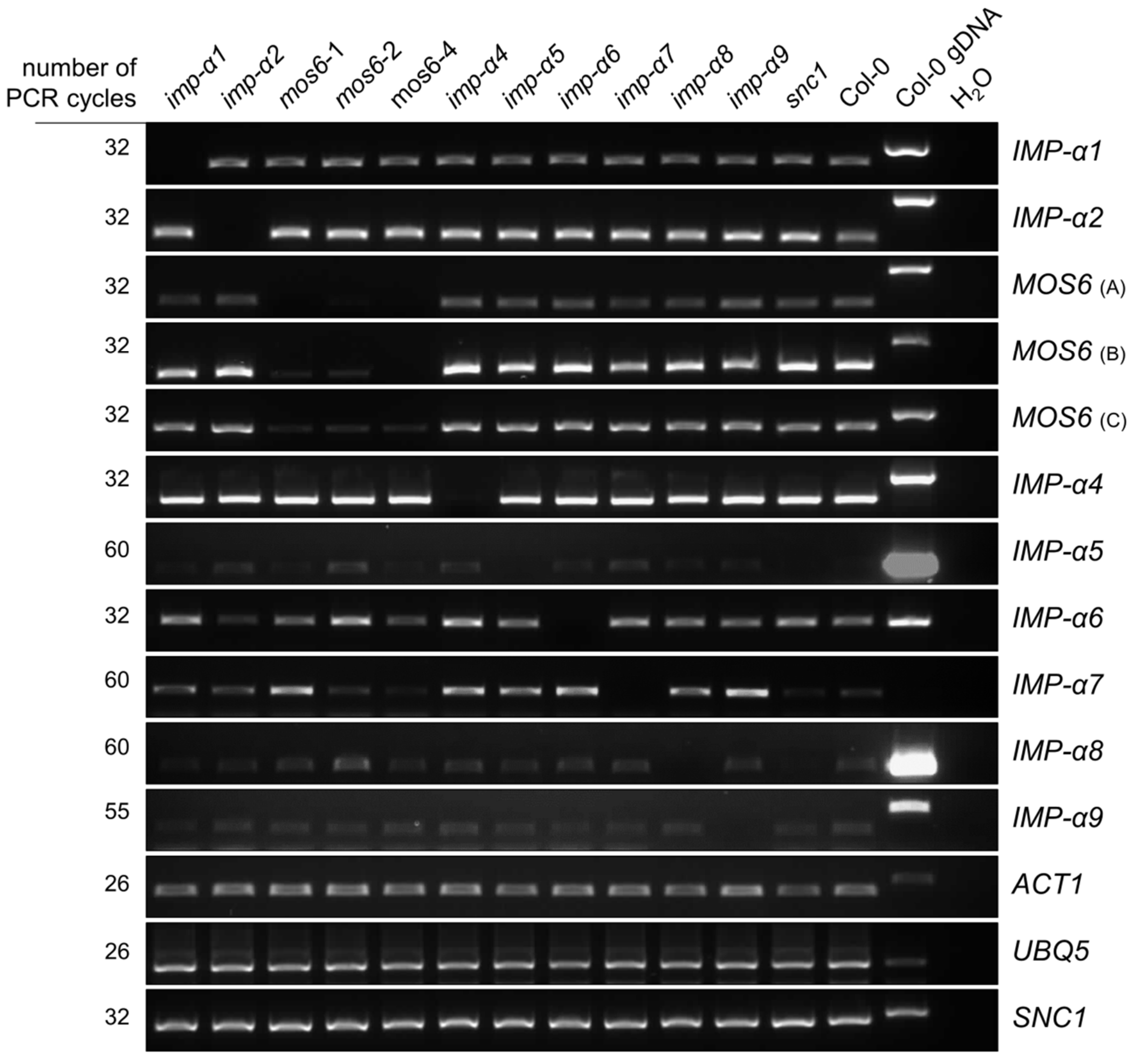


**Figure S3.** *Imp-*$\alpha$ single mutants do not show obviously altered expression of the remaining functional *IMP-*$\alpha$ genes or *SNC1*. RT-PCRs were performed with the indicated number of PCR cycles, using primers for the nine *Arabidopsis* $\alpha$*-IMPs* indicated in Figure S1 and cDNA transcribed from RNA of four week old plants of the indicated genotypes. *ACTIN1* (*ACT1*) and *UBIQUITIN5* (*UBQ5*) were used as controls. PCR products were separated by agarose gel electrophoresis and stained by ethidium bromide.

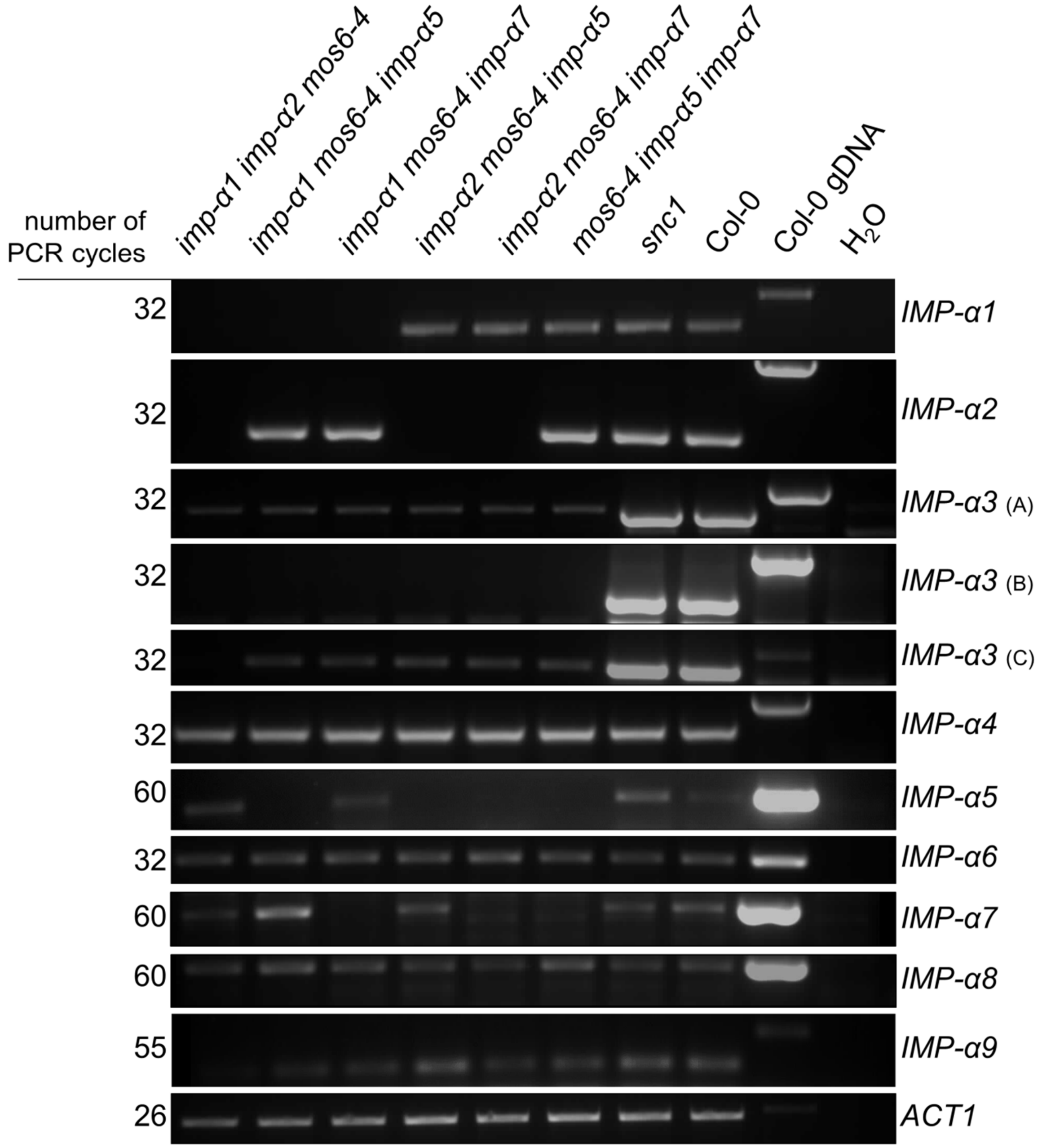


**Figure S4.** *Imp-$\alpha$* triple mutants or *snc1* mutants do not show obviously altered expression of the remaining functional *IMP-$\alpha$´s*. RT-PCRs were performed with the indicated number of PCR cycles, using primers indicated in Figure S1 and cDNA transcribed from RNA of four week old plants of the indicated genotypes. *ACTIN1* (*ACT1*) was used as a control. PCR products were separated by agarose gel electrophoresis and stained by ethidium bromide.

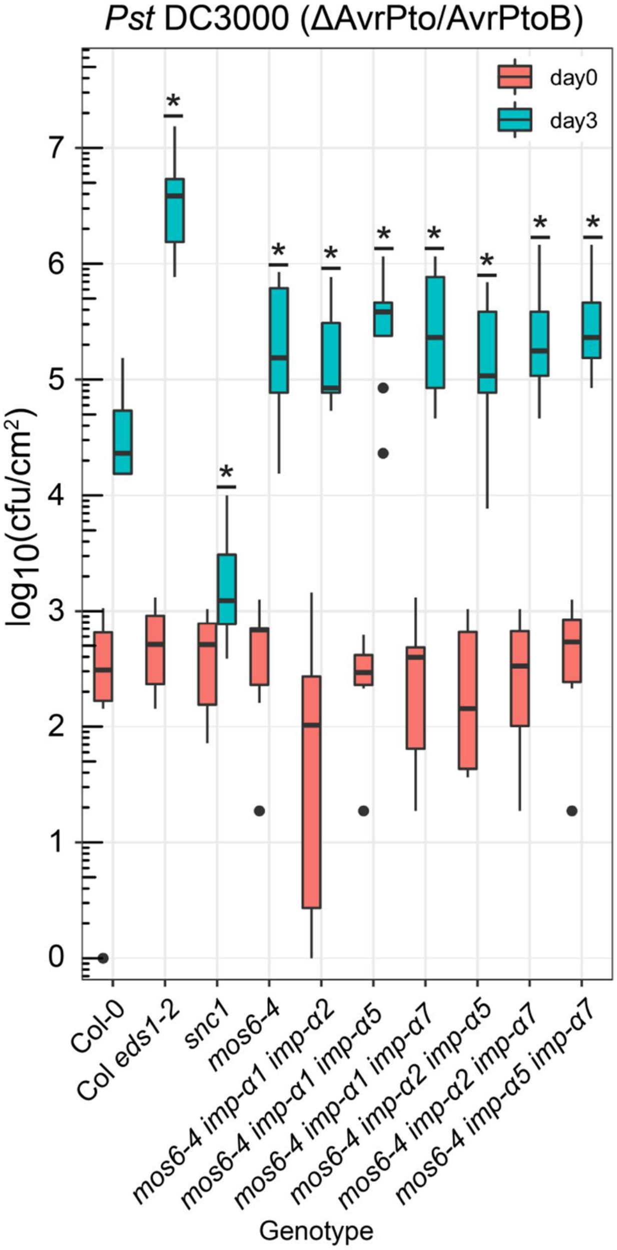


**Figure S5.** Only the *mos6* allele contributes to immunity against mildly virulent *Pst* in *imp-α* triple mutant lines. Four week old plants of the indicated genotypes were vacuum infiltrated with a *Pst* DC3000 (ΔAvrPto/AvrPtoB) suspension of $1 \times 10^5$ cfu $ml^{-1}$. Colony-forming units (cfu) within the infiltrated plant tissues were quantified immediately (day 0) or three days after infiltration (day 3). Data is presented as boxplots (day 0: n=6; day 3: n=9), outliers are indicated as black dots, underlined asterisks indicate statistically significant differences to Col-0 (one-way ANOVA; Tukey's test, $P < 0.05$). The experiment was repeated three times with similar results.

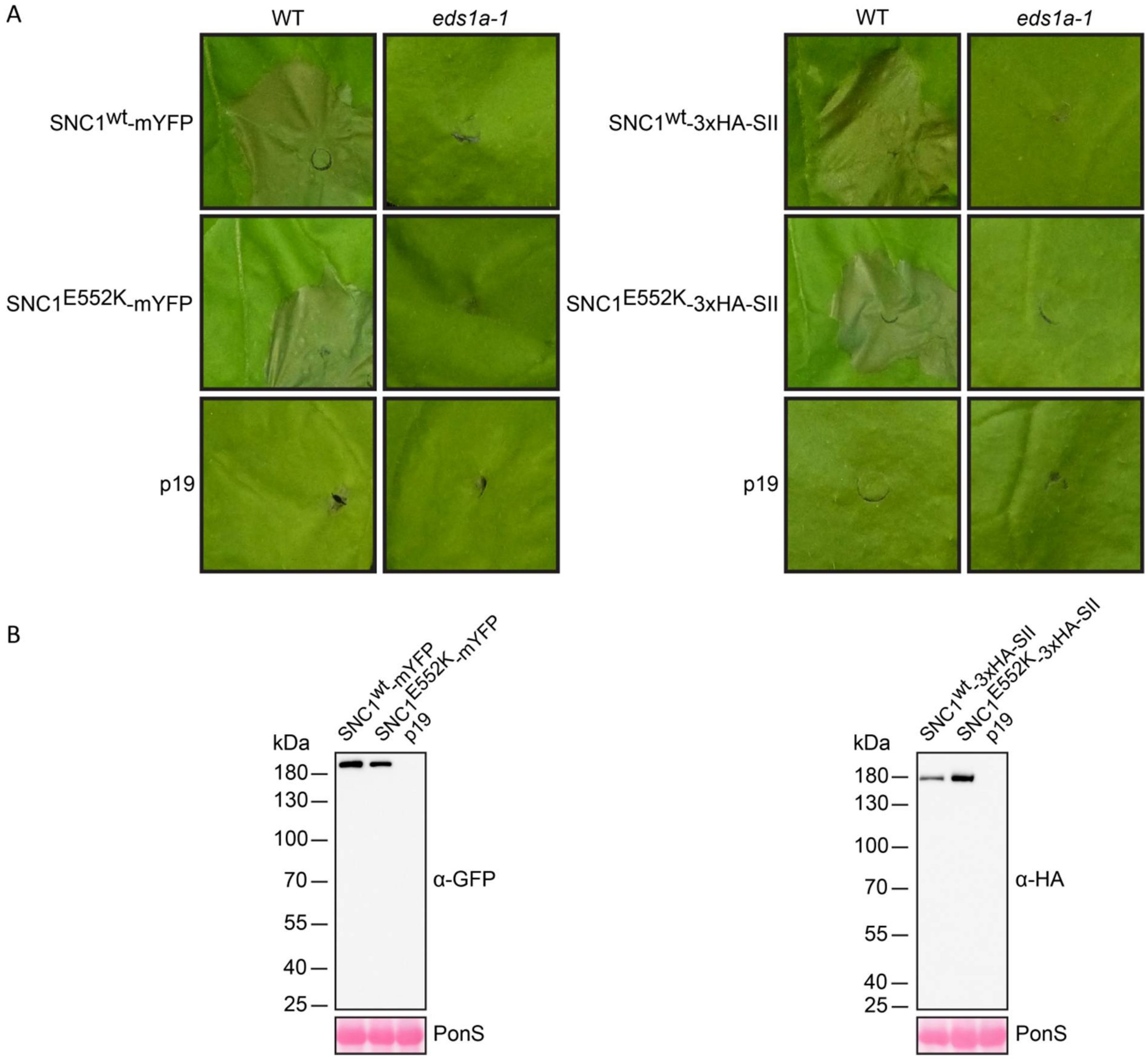


**Figure S6.** SNC1$^{wt}$ and SNC1$^{E552K}$ can be transiently expressed to detectable levels in the *Nicotiana benthamiana eds1a-1* mutant without induction of a cell death response. (A) *Agrobacterium*-mediated transient expression of mYFP-tagged or 3xHA-StrepII (3xHA-SII)-tagged SNC1$^{wt}$ and SNC1$^{E552K}$ in *Nicotiana benthamiana* wildtype or *eds1a-1* mutant plants, together with the p19 silencing suppressor. Cell death was visualized 4 d after infiltration (4 dpi) of *Agrobacteria*. p19 only expression was used as a control. (B) Immunoblots of total protein extracts derived from *Nicotiana benthamiana eds1a-1* plants, infiltrated with *Agrobacteria* for transient expression of mYFP-tagged or 3xHA-SII-tagged SNC1$^{wt}$ and SNC1$^{E552K}$. Total extracts were taken from infiltrated areas of *Nbeds1a-1* plants 4 dpi and separated on 10 % SDS polyacrylamide gels, blotted onto nitrocellulose membranes and probed with $\alpha$-GFP or $\alpha$-HA primary antibodies. Ponceau S (PonS) staining of the membrane was used to monitor loading.

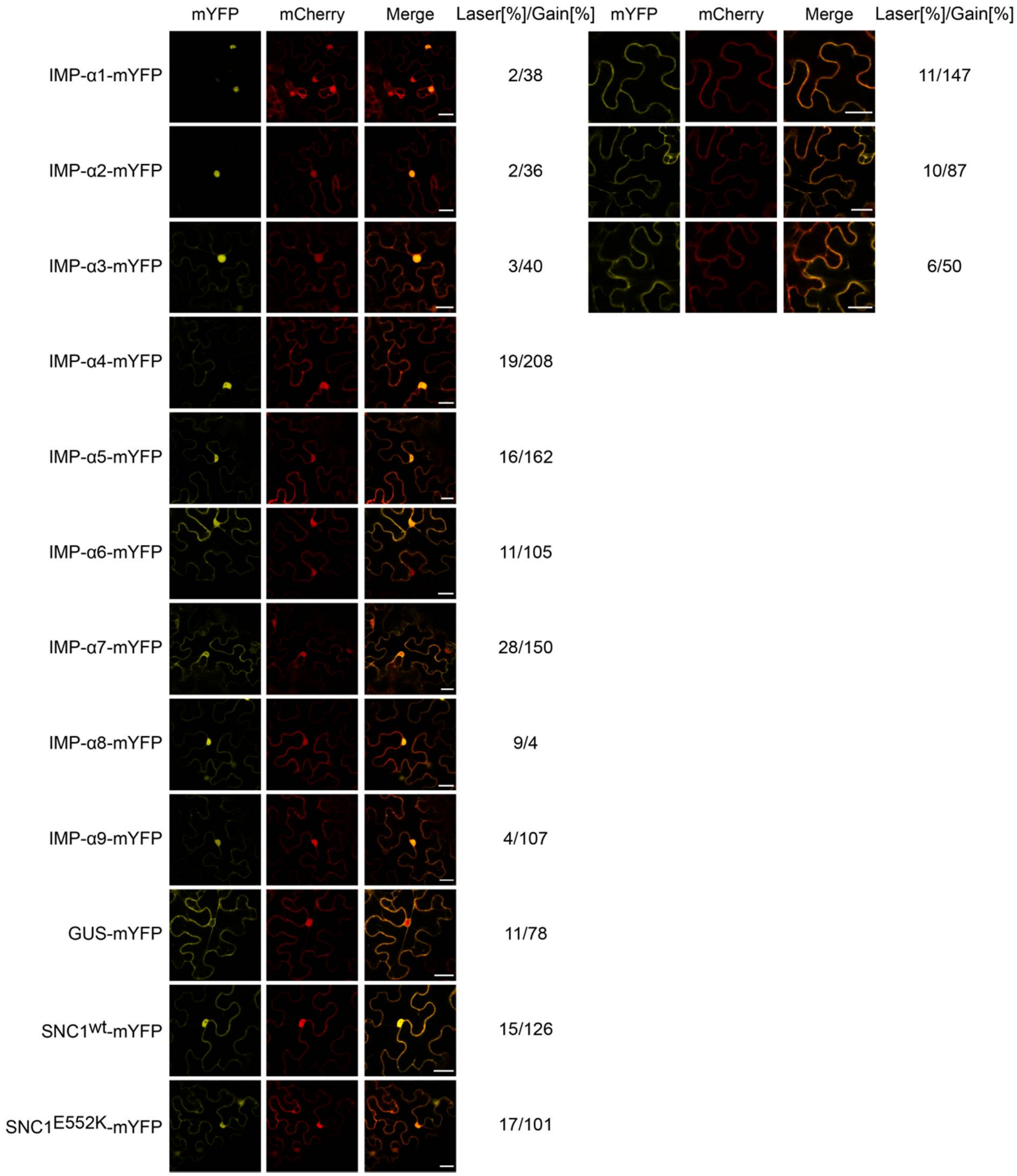


**Figure S7.** Transiently expressed α-IMPORTINS, $SNC1^{wt}$ and $SNC1^{E552K}$ show a nuclear-cytoplasmic localization in *Nicotiana benthamiana*. Confocal laser scanning microscopy images of *Nicotiana benthamiana eds1a-1* plants, transiently co-expressing free mCherry together with mYFP-tagged α-IMPORTINS, $SNC1^{wt}$ and $SNC1^{E552K}$, respectively, 2 days after *Agrobacterium* infiltration. GUS-mYFP expression was used as control. All constructs were expressed together with the silencing suppressor p19. mYFP fluorescence is shown in yellow and mCherry fluorescence is shown in red. Respective laser power and detector gain are indicated. Sections defocusing the nuclei are shown in addition for IMP-α1, IMP-α2 and IMP-α3 to visualize cytoplasmic localization. Scale bars = 25 µm.

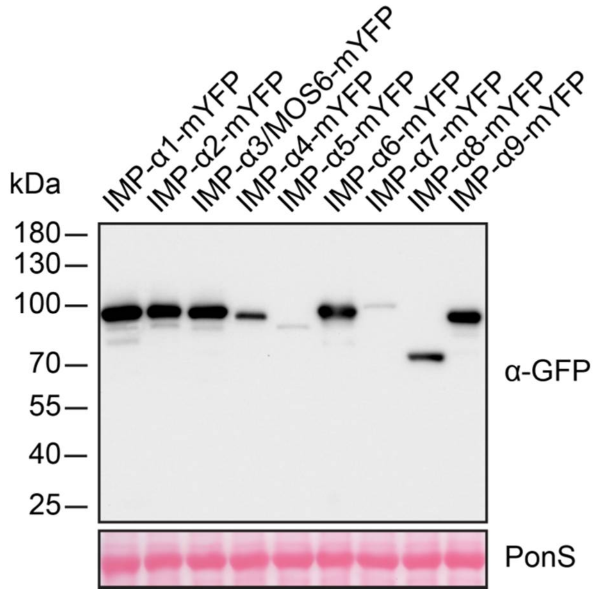


**Figure S8.** Accumulation of full length mYFP-tagged IMPORTIN-$\alpha$ proteins upon transient expression in *N. benthamiana*. Immunoblot analysis of total protein extracts derived from *Nicotiana benthamiana eds1a-1* plants, transiently co-expressing free mCherry together with mYFP-tagged $\alpha$-IMPORTINS for confocal laser scanning microscopy, 2 days after *Agrobacterium* infiltration (Figure S7). Total extracts were taken from infiltrated areas 2 dpi and were separated on 10 % SDS polyacrylamide gels, blotted onto nitrocellulose membranes and probed with $\alpha$-GFP primary antibody for IMP-$\alpha$-mYFP detection. Ponceau S (PonS) staining of the membrane was used to monitor loading.

**Table S1.** NLS predictions for SNC1. NLS sequences and their locations within the SNC1 protein sequence are shown. A low cut-off value (0.5) for two-state Hidden Markov Model static prediction was used in NLStradamus. The respective scores for NLS predictions in NLSmapper are indicated.

| | **NLStradamus** | **NLSmapper (score)** |
|---|---|---|
| 1 | | $_{17}$GSRRYDVFPSFRGEDVRDSFLSHLLKELRGK$_{47}$ (2.8) |
| 2 | $_{497}$AKSKGNPGKR$_{506}$ | $_{502}$NPGKRRFLTNF$_{512}$ (2.5) |
| 3 | | $_{1360}$RSEKRMRMT$_{1368}$ (3.0) |

**Table S2.** Primers used in this study. The name, sequence and use of each primer is indicated.

| Name | Sequence 5´-3´ | Use |
|---|---|---|
| IMP-α6_CRISPR_Fwd | GATTGAGACTTACAATTGGAGGCGA | sgRNA/target site |
| IMP-α6_CRISPR_Rev | AAACTCGCCTCCAATTGTAAGTCTC | sgRNA/target site |
| IMP-α1_Fwd (RP) | GCGTGATCAATAGTGGTGC | RT-PCR/genotyping |
| IMP-α1_Rev (LP) | GAGAACCATCAACACCTGGAG | RT-PCR/genotyping |
| IMP-α2_Fwd (LP) | AATCTGTCATTGAGGCAGGTG | RT-PCR/genotyping |
| IMP-α2_Rev (RP) | TCAAACCAGCTTCACACACAG | RT-PCR/genotyping |
| mos6-1/IMP-α3_A_Fwd | CATTAATAACGAACCTGCCAC | genotyping |
| mos6-1/IMP-α3_A_Rev | TGCTAGAACCAAATTGCAGC | genotyping |
| mos6-2/IMP-α3_B_Fwd | TATCTGATCTGCATTTCCAGC | genotyping |
| mos6-2/IMP-α3_B_Rev | CTCCTAAGTACACGCTCTTG | genotyping |
| mos6-4/IMP-α3_C_Fwd (RP) | TCATTCGTCGCCATCAGTGC | RT-PCR/genotyping |
| mos6-4/IMP-α3_C_Rev (LP) | GAACATTGGTGCTTCCGAAC | RT-PCR/genotyping/ |
| IMP-α4_Fwd (LP) | GCTGCATGGGCTTTGAC | RT-PCR/genotyping |
| IMP-α4_Rev (RP) | GCAAGCATCAGTGAGAACCTC | RT-PCR/genotyping/ |
| IMP-α5_LP | CTTCGCCGGAGAAAACACTAC | genotyping |
| IMP-α5_RP | AACAAAGAGTGGCACAACACC | genotyping |
| IMP-α6-1_LP | GGCAATCTCTAATGCAACTTCTGG | genotyping |
| IMP-α6-1_RP | GACATGTAGATGTTGATTCTGC | genotyping |
| IMP-α6-2_LP | ATGTCTTACAAACCAAGCGCGAAG | genotyping |
| IMP-α6-2_RP | CCAATTGTACGGAGAGCTGGAATCA | genotyping |
| IMP-α7_LP | GCCATAGCTGGAGGCTCTTAC | genotyping |
| IMP-α7_RP | AAAACTCATTAAACCCAGCCG | genotyping |
| IMP-α8_LP | CTTCTCCAGTGGTGCTTGTTC | genotyping |
| IMP-α8_RP | TCTTCAAACCAATTCGTCACC | genotyping |
| IMP-α9_LP | AAGCTGCTAGGCTTGGTCTTC | genotyping |
| IMP-α9_RP | TCATGACGTCGAAACCCTAAC | genotyping |
| snc1_LP | TGGTTTTGAAGTCAGTTACG | genotyping |
| snc1_RP | CAAGTTGAGATCGGTTGG | genotyping |
| mos6-1/IMP-α3_A_Fwd | CTCAGACCTAGCGCGAAGAC | RT-PCR |
| mos6-1/IMP-α3_A_Rev | TCCAGCAACCATAGCAGGTAG | RT-PCR |
| mos6-2/IMP-α3_B_Fwd | CTAGTGAAGATGTCCGCGAAC | RT-PCR |
| mos6-2/IMP-α3_B_Rev | AAGCGCTTGCTGGTCG | RT-PCR |
| IMP-α5_Fwd | ATACTTGGTGGAGCAGAATTGC | RT-PCR |
| IMP-α5_Rev | CAACCATCACCACCTTCATCT | RT-PCR |
| IMP-α6_Fwd | GTTTCTCGTGAGCCAAGGC | RT-PCR |
| IMP-α6_Rev | ACCAAAGTTGAATCCACCCG | RT-PCR |
| IMP-α7_Fwd | TGCATCAAACCGTTGTGCGA | RT-PCR |
| IMP-α7_Rev | AGGTCCGCAGTGCATCTC | RT-PCR |
| IMP-α8_Fwd | ATACATGGCGGAGCAGAGTT | RT-PCR |
| IMP-α8_Rev | CACCTGAAAGTCCACATCATCAC | RT-PCR |
| IMP-α9_Fwd | AAGCTGCTAGGCTTGGTCTTC | RT-PCR |
| IMP-α9_Rev | TTCATCGATTCCATAATCTTCACC | RT-PCR |
| ACTIN1_RT-PCR_Fwd | CGATGAAGCTCAATCCAAACGA | RT-PCR |

| | | |
|---|---|---|
| ACTIN1_RT-PCR_Rev | CAGAGTCGAGCACAATACCG | RT-PCR |
| UBQ5_RT-PCR_Fwd | GACGCTTCATCTCGTCC | RT-PCR |
| UBQ5_RT-PCR_Rev | GTAAACGTAGGTGAGTCCA | RT-PCR |
| SNC1_RT-PCR_Fwd | CATTTTCAGACTTACAAGACTTGAGC | RT-PCR |
| SNC1_RT-PCR_Rev | GTAAGTTGATATCTTCTTCAGATGTCC | RT-PCR |
| gIMP-α1.D-TOPO_Fwd | CACCATGTCACTGAGACCCAACG | cloning |
| gIMP-α1.D-TOPO_Rev.Δstop | GCTGAAGTTGAATCCTCCG | cloning |
| gIMP-α2.D-TOPO_Fwd | CACCATGTCTTTGAGACCTAACGC | cloning |
| gIMP-α2.D-TOPO.Rev_Δstop | CTGGAAGTTGAATCCACCTG | cloning |
| gIMP-α4.D-TOPO_Fwd | CACCATGTCGCTGAGGCCGAG | cloning |
| gIMP-α4.D-TOPO_Rev.Δstop | GGCAAATTTGAATCCACCAACG | cloning |
| gIMP-α5.D-TOPO_Fwd | CACCATGTCCTTGCGACCGAGC | cloning |
| gIMP-α5.D-TOPO_Rev.Δstop | ACGAGAAAAATCAAACTGGAATTCC | cloning |
| gIMP-α7.D-TOPO_Fwd | CACCATGAAGGGAGGAGAGACAATG | cloning |
| gIMP-α7.D-TOPO_Rev.Δstop | AGGTCCGCAGTGCATCTC | cloning |
| gIMP-α8.D-TOPO_Fwd | CACCATGGCTTGGAAAACAGAG | cloning |
| gIMP-α8.D-TOPO_Rev.Δstop | CACCTGAAAGTCCACATCATC | cloning |
| gIMP-α9.D-TOPO_Fwd | CACCATGGCGGATGATGGCTC | cloning |
| gIMP-α9.D-TOPO_Rev.Δstop | TTCATCGATTCCATAATCTTCACC | cloning |
| gSNC1.D-TOPO_Fwd | CACCATGGAGATAGCTTCTTCTTCTGGC | cloning |
| gSNC1.D-TOPO_Rev.Δstop | GTTACCAGAAACAGGAAACAAGATAGG | cloning |
